# A master regulatory loop that activates genes in a temporally coordinated manner in muscle cells of ascidian embryos

**DOI:** 10.1101/2024.08.27.610013

**Authors:** Izumi Oda, Yutaka Satou

## Abstract

Ascidian larval muscle cells present a classic example of autonomous development. A regulatory mechanism for these cells has been extensively investigated, and the regulatory gene circuit has been documented from maternal factors to a muscle specific gene. In the present study, we comprehensively identified genes expressed specifically in ascidian muscle cells, and found that all of them are under control of a positive regulatory loop of *Tbx6-r.b* and *Mrf*, the core circuit identified previously. We also found that several transcription factors under control of the *Tbx6-r.b*/*Mrf* regulatory loop resulted in various temporal expression profiles, which are probably important for creating functional muscle cells. These results, together with results of previous studies, provide an exhaustive view of the regulatory system enabling autonomous development of ascidian larval muscle cells. It shows that the *Tbx6-r.b*/*Mrf* regulatory loop, but not a single gene, serves a “master” regulatory function. This master regulatory loop not only controls spatial gene expression patterns, but also governs temporal expression patterns in ascidian muscle cells.

## Introduction

Tadpole larvae of ascidians, which belong to the sister group of vertebrates, differentiate 36 muscle cells in their tails. The anterior 28 muscle cells (B-line muscle cells) have been investigated as a classical example of autonomous development. The core circuit of the genetic pathway behind this autonomous differentiation is a feedback loop consisting of two transcription factor genes, *Tbx6-r.b* and *Mrf* (Imai et al., 2006; Meedel et al., 2007; Yagi et al., 2005; Yu et al., 2019).

*Mrf* is the sole ortholog for the vertebrate myogenic factor genes, *MyoD*, *Myf5*, *Myogenin*, and *Mrf4* (Araki et al., 1994; Meedel et al., 1997). Among these myogenic transcription factors, MyoD was first identified as a factor that can convert mouse fibroblast cells to myoblast cells (Lassar et al., 1986). This factor has been regarded as a “master control gene”, positioned at the top of a genetic pathway specifying muscle fate. In fact, knockout of myogenic factor genes greatly impairs myogenesis in vertebrate embryos (Hasty et al., 1993; Kassar-Duchossoy et al., 2004; Nabeshima et al., 1993; Rudnicki et al., 1993). On the other hand, the myogenetic factor gene plays a relatively minor role in differentiation of muscle of flies, nematodes, and sea urchin (Balagopalan et al., 2001; Chen et al., 1994; Venuti et al., 1993). Similarly, knockdown of *Mrf* in ascidian embryos using specific morpholino oligonucleotides does not completely eliminate muscle cells (Imai et al., 2006; Meedel et al., 2007). Indeed, *Myl.c*, which encode myosin light chain, is expressed in early embryos independently of *Mrf*, but under control of *Tbx6-r.b* (Yu et al., 2019). On the other hand, as mentioned above, *Mrf* together with *Tbx6-r.b*, constitutes a positive feedback loop, and many muscle-structural genes may be under control of this feedback loop. The first question in the present study is whether this regulatory loop has a “master regulatory” role. In other words, are all genes specifically expressed in the muscle lineage regulated by this feedback loop?

The activity of this feedback loop subsides gradually, and *Tbx6-r.b* expression mostly disappears before the tailbud stage (Takatori et al., 2004). Instead of *Tbx6-r.b*, *Tbx15/18/22* begins to be expressed under control of the regulatory loop and maintains expression of *Myl.c* in late embryos and larvae through a shared binding site for Tbx6-r.b and Tbx15/18/22 in the upstream regulatory region of *Myl.c*. In addition to this shared binding site, *Myl.c* has a site that binds Tbx6-r.b, but not Tbx15/18/22, and *Myl.c* is activated through this site before gastrulation. That is, *Myl.c* is regulated by *Tbx6-r.b* in early embryos, then by *Tbx6-r.b* and *Mrf*, and finally by *Tbx15/18/22* (Yu et al., 2019). This observation indicates that temporal expression patterns of genes expressed in muscle cells can be regulated temporally by a combination of transcription factors. A previous comprehensive gene expression assay showed various temporal expression patterns of muscle-specific genes (Satou et al., 2001). On the basis of these observations, the next question is how expression of muscle-specific genes is temporally regulated. *Snai*, *Meox* (*Mox*), and *Tcf15-r* (*Paraxis-like*), which encode transcription factors, are expressed in the muscle lineage (Cao et al., 2019; Fujiwara et al., 1998; Imai et al., 2004; Satou et al., 2001); therefore, they may contribute to temporal regulation of muscle-specific genes as *Tbx15/18/22* does.

Together with results of previous studies, answers to these questions will uncover the entire genetic pathway from maternal factors to expression of muscle-specific genes through the lifetime of muscle cells of ascidian larvae, and will illuminate how the gene regulatory system coordinates gene expression to differentiate functional muscle cells.

## Results

### All muscle-specific genes are regulated under control of *Tbx6-r.b* and *Mrf*

First, we used publicly available single-cell transcriptome data for late tailbud embryos to identify muscle-specific genes (Cao et al., 2019). We mapped the raw data to a gene model set (KY21 set) that was recently published (Satou et al., 2022). We found that 275 genes were significantly enriched in a cluster representing muscle cells (p<0.01; mean UMI count>1), but this set included genes that are expressed not only in muscle, but also in other tissues. To exclude such genes, we calculated mean expression counts per gene per cluster, and found that expression counts of 102 genes in the muscle cluster were ≥10 times as abundant as those in all other clusters. This set contained nine genes encoding muscle actin, six genes encoding myosin heavy chain, six genes encoding myosin light chain, one gene encoding troponin I, one gene encoding troponin T, two genes encoding troponin C, two genes encoding tropomyosin, and one gene encoding muscle-type creatine kinase; thus, muscle-specific genes were successfully identified with this method. In addition, we randomly chose six genes from this list, and confirmed that these genes are expressed specifically in muscle cells (Figure S1). On the other hand, we excluded one gene (KY21.Chr1.1843) from the list, because this gene was a single-exon gene encoded in an intron of another gene. Specifically, it encoded a short protein (30 amino acids) with no clear similarity to proteins of other animals. Moreover, its GC content (25%) was much lower than the average (35%) over the genome (Dehal et al., 2002), and this low GC content potentially causes mis-mapping of sequencing tags and makes downstream assays difficult. Thus, in subsequent analyses, we used the remaining 101 genes (Table S1).

We then performed RNA-sequencing of tailbud embryos injected with a control morpholino oligonucleotide (MO), a MO against *Tbx6-r.b* or a MO against *Mrf*. We took late tailbud embryos for both morphants, and we also took middle tailbud embryos for *Tbx6-r.b* morphants, because *Tbx6-r.b* expression ceases earlier than *Mrf* expression (Yu et al., 2019). While the *Mrf* MO has been used in previous studies (Imai et al., 2006; Yu et al., 2019), the *Tbx6-r.b* MO used in the present study was different from the MO we used previously. So we first examined expression of two known targets [muscle actin genes and *Myl.c* (Yagi et al., 2005)] of *Tbx6-r.b* in embryos injected with the new MO, and found that this new MO downregulated expression of these two genes in the same way as the old one, while expression levels of control genes (*Cdk5* and *Slk/Stk10*) hardly changed (Figure S2).

As shown in Figure 1A, the above 101 genes were all significantly down-regulated in *Tbx6-r.b* morphant embryos and/or *Mrf* morphant embryos (adjusted p<0.01; Figure 1A; Table S2). Most of them (69 genes) were downregulated in all three experiments (Figure 1B). This is not surprising because *Tbx6-r.b* and *Mrf* constitute a positive feedback loop (Yu et al., 2019). As further confirmation, we performed an RNA-seq analysis using late tailbud embryos injected with morpholino oligonucleotides of *Tbx6-r.b* and *Mrf* simultaneously, and found that 99 genes were downregulated significantly (adjusted p-value <0.01; Figure 1A; Table S2). Expression of the remaining two genes (*Slc38a3* and *Obscn*) was also downregulated with modest adjusted-p-values (0.017 and 0.024). We quantified their mRNA levels in double morphants of *Tbx6-r.b* and *Mrf* with reverse transcription and quantitative PCR (RT-qPCR), and found that these genes were indeed downregulated in double morphants of *Tbx6-r.b* and *Mrf* (Figure S3). These data indicate that the 101 muscle-specific genes are under control of the regulatory loop of *Tbx6-r.b* and *Mrf*.

**Figure 1.**
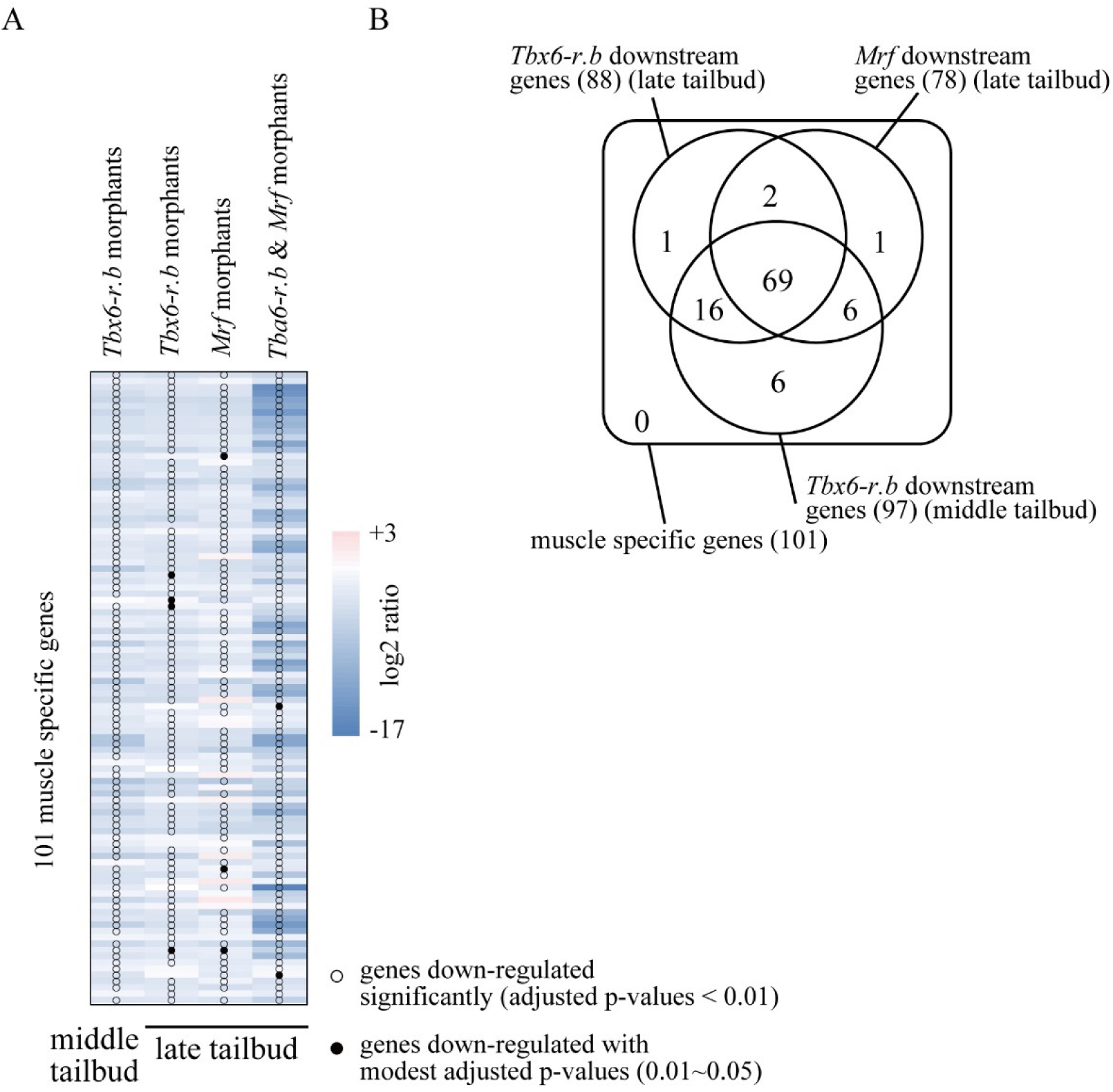
RNA-sequencing experiments revealed that all 101 muscle-specific genes are under control of the regulatory loop of *Tbx6-r.b* and *Mrf*. (A) A heatmap of changes in expression of the 101 muscle-specific genes between control and morphant embryos designated above. RNA-sequencing results were analyzed with Deseq2 (Love et al., 2014). Genes, expression of which was significantly downregulated, are shown by dots. (B) A Venn diagram that shows overlaps among three RNA-sequencing experiments (threshold p-value, 0.01).

### Temporal expression profiles of genes specifically expressed in the muscle lineage

To examine temporal expression profiles of the above 101 muscle-specific genes, we used publicly available RNA-sequencing data that quantified RNA-levels at various stages (Brozovic et al., 2018). We mapped data of early gastrula, middle gastrula, middle neurula, middle tailbud, and larval stages to the KY21 gene model set, and found that 51 genes are most abundantly expressed in larvae (Lv-peak genes), 41 genes in middle tailbud embryos (mTB-peak genes), 7 genes in middle neurula embryos (mN-peak genes), and 2 genes in middle gastrula embryos (mG-peak genes) (Figure 2A; Table S3).

**Figure 2.**
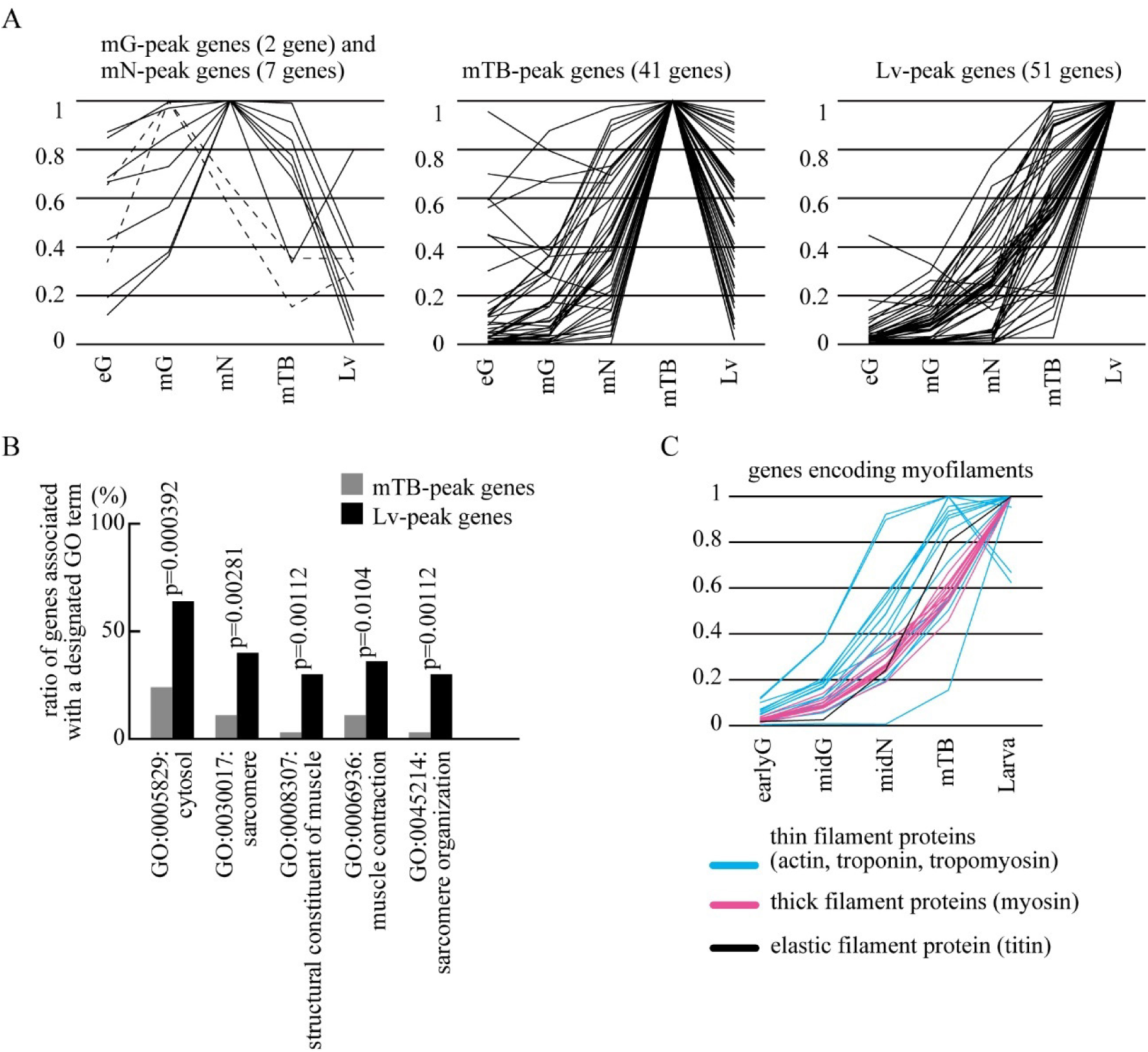
Genes encoding myofiber proteins tend to be expressed abundantly in larvae. (A) Temporal expression patterns of the 101 muscle-specific genes, which were calculated using publicly available RNA-sequencing data (Brozovic et al., 2018). TPM values were converted to relative values against the highest value for each gene. Genes with peak expression at the middle gastrula (mG, broken lines) or middle neurula (mN, solid lines) stages are shown in the left panel (mG-peak genes and mN-peak genes). Genes with peak expression at the middle tailbud (mTB) and larval (Lv) stages are shown in the middle and right panels (mTB-peak genes and Lv-peak genes). No genes showed peak expression at the early gastrula (eG) stage. (B) mTB-peak genes and Lv-peak genes were analyzed using gene ontology terms (Ashburner et al., 2000). Differences between the two populations were examined using Fisher’s exact tests. GO terms that passed the multiple testing correction using the Benjamini-Hochberg method (threshold value = 0.1) are shown with p-values of Fisher’s exact tests. (C) Temporal expression profiles of genes encoding myofiber proteins.

Among the 101 genes, 94 encoded proteins with clear homologs in the human proteome (BLASTP E-value, ≤1×10^-5^). Using gene ontology (GO) terms associated with these homologous human proteins, we analyzed differences in GO terms between the mTB-peak genes (n=38) and Lv-peak genes (n=47). The remaining two gene groups were not analyzed because gene numbers were too small for statistical comparisons. Eight cellular-component GO-terms, four molecular-function GO-terms, and two biological-process GO-terms were associated with 15 or more proteins (Table S4; Table S5).

Among the eight cellular-component GO-terms, “cytosol” and “sarcomere” were significantly more frequently associated with Lv-peak genes than mTB-peak genes (Fisher’s exact test p-values, 3.92×10^-4^, and 2.81×10^-3^). Accordingly, one molecular function GO term, “structural constituent of muscle” (p-value, 1.12×10^-3^), and two biological process terms, “muscle contraction” (p-value=1.04×10^-2^) and “sarcomere organization” (p-value, 1.12×10^-3^), were significantly more frequently associated with Lv-peak genes (Figure 2B; Table S5). Consistently, 25 of 28 genes encoding myofilament proteins (myosin, actin, tropomyosin, troponin, and titin) were expressed most abundantly at the larval stage (Figure 2C). On the other hand, “plasma membrane” was more frequently associated with mTB-peak genes (P-value, 2.44×10^-2^), although multiple corrections using the Benjamini-Hochberg method (threshold=0.1) did not result in statistical significance. Indeed, two genes encoding cholinergic receptors (KY21.Chr7.717 and KY21.Chr7.961) and three of four genes encoding solute carrier proteins (KY21.Chr1.1444, KY21.Chr3.8, and KY21.Chr6.592) were most abundant at the middle tailbud stage (Table S5). These observations suggest that populations of mTB-peak genes and Lv-peak genes are qualitatively different, and that genes expressed in muscle cells are temporally regulated according to protein functions.

### Temporal expression profiles of transcription factor genes in the muscle lineage

A mechanism that temporally regulates muscle-specific genes has partly been revealed. *Myl.c*, which encodes a myosin light chain, has two cis-regulatory elements. The first of these contains a Tbx6-r.b-binding site and regulates expression in early embryos; *Tbx6-r.b* initiates expression earlier than *Mrf*. On the other hand, the second contains binding sites for Tbx6-r.b and Mrf and regulates expression in late embryos (Yu et al., 2019). The second also binds Tbx15/18/22, which is expressed in late embryos, and maintains expression in late embryos and larvae (Yu et al., 2021). In addition, *Meox* (*Mox*), *Tcf15-r* (*Paraxis-like*), and *Snail* are expressed in the muscle lineage (Cao et al., 2019; Fujiwara et al., 1998; Imai et al., 2004; Satou et al., 2001). *Meox* expression becomes detectable specifically in the muscle lineage by *in situ* hybridization at the late gastrula stage, and continues to be expressed at the middle tailbud stage (Imai et al., 2004) (Figure S4). *Tcf15-r* (*Paraxis-like*) is also expressed specifically in the muscle lineage from late gastrula to tailbud embryos (Satou et al., 2001) (Figure S4). *Snail* expression in the muscle lineage begins at the 32-cell stage, and is hardly detectable before the middle tailbud stage (Fujiwara et al., 1998; Imai et al., 2004).

To more precisely understand temporal expression profiles of these transcription factor genes, which are expressed in the muscle lineage, we first examined expression profiles of these transcription-factor genes by reverse-transcription and quantitative PCR (RT-qPCR). As we showed previously (Yu et al., 2021), *Tbx6-r.b* expression was decreased and hardly undetectable before the late tailbud stage, whereas expression of *Mrf* and *Tbx15/18/22* was readily detected in larvae (Figure 3A–C). *Snai* expression was detected at the middle tailbud stage, but rapidly decreased before the late tailbud stage (Figure 3D). Expression profiles of *Meox* and *Tcf15-r* were similar. These genes were expressed most abundantly at the tailbud stage, and were barely detectable in larvae (Figure 3EF). These observations indicate the possibility that *Meox*, *Tcf15-r*, and *Snail* are involved in temporal regulation of muscle structural genes as in the case of *Tbx15/18/22*.

**Figure 3.**
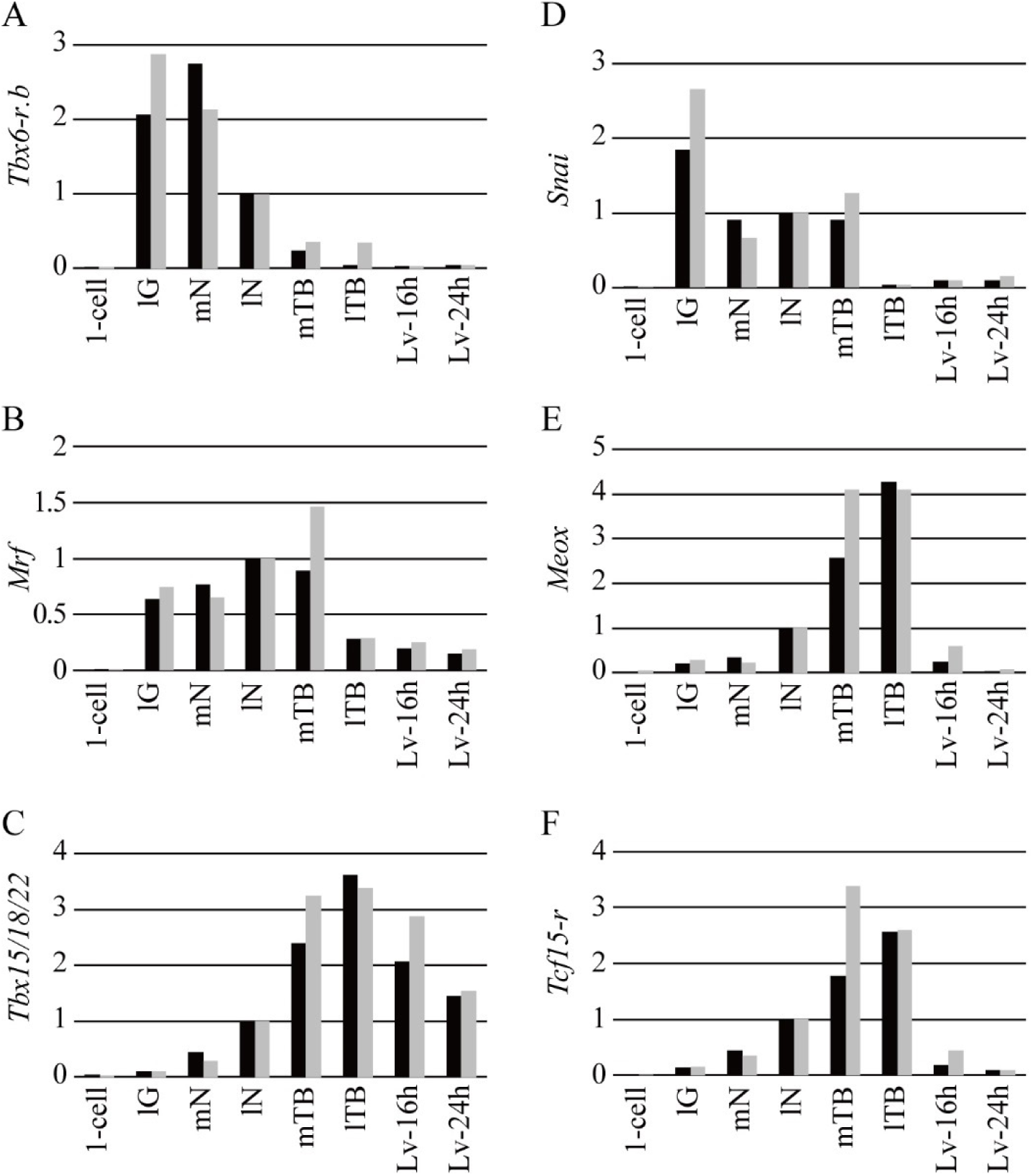
Temporal gene expression profiles of six transcription factor genes. Expression levels were quantified by RT-qPCR assays at 1-cell, late gastrula (lG), middle neurula (mN), late neurula (lN), middle tailbud (mTB), late tailbud II (lTB), 16h-larva (Lv-16h), and 24h-larva stages (Lv-24h), and calculated as relative values against expression levels at the late neurula stage. Results from two batches of embryos are represented with different colors.

### *Meox* and *Tcf15-r* downstream genes are highly expressed in larvae

Because the above RNA-seq experiments showed that *Meox* and *Tcf15-r* were down-regulated in double morphants of *Tbx6-r.b* and *Mrf* (Figure 4A; Table S2), *Meox* and *Tcf15-r* may regulate gene expression in late embryos under control of the regulatory loop of *Tbx6-r.b* and *Mrf*. To test this hypothesis, we first performed RNA-seq experiments using late tailbud embryos developed from eggs injected with a MO against *Meox* or *Tcf15-r*. We took biological duplicates for each experiment and found that six and nine genes were downregulated significantly in *Meox* and *Tcf15-r* morphants, respectively (Figure 4B; Table S2). To compare these results with our previous result for *Tbx15/18/22*, we reanalyzed *Tbx15/18/22* data using the latest gene model set (Satou et al., 2022), which we used in the present study. There were overlaps among downstream genes of these three factors (Figure 4C).

**Figure 4.**
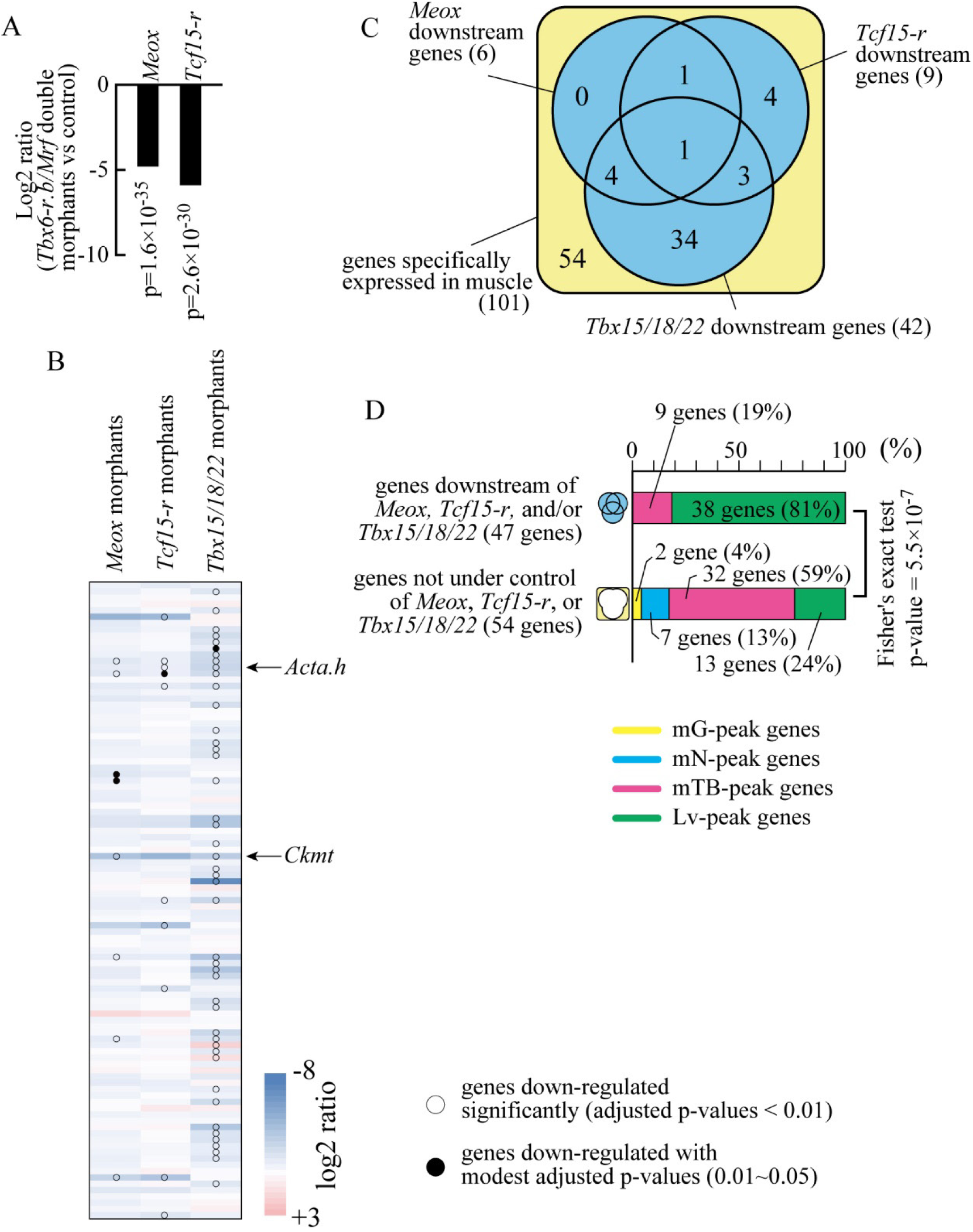
*Meox* and *Tcf15-r* regulate subsets of muscle-specific genes, most of which are expressed most abundantly in larvae. (A) RNA-sequencing experiments using double morphants of *Tbx6-r.b* and *Mrf* (see Figure 1) showed that *Meox* and *Tcf15-r* are under control of *Tbx6-r.b* and *Mrf*. Adjusted p-values, which are shown in the panel, were calculated using Deseq2 (Love et al., 2014). (B) A heatmap of changes in expression of the 101 muscle-specific genes between control and morphant embryos of *Meox*, *Tcf15-r*, or *Tbx15/18/22*. RNA-sequencing results were analyzed with Deseq2 (Love et al., 2014). Genes, expression of which was significantly downregulated, are shown with dots. We used RNA-sequencing data for *Tbx15/18/22* morphants that were published previously (Yu et al., 2021) and re-mapped them to the latest gene model set (Satou et al., 2022). (C) A Venn diagram that shows overlaps among genes under control of *Meox*, *Tcf15-r*, and/or *Tbx15/18/22*. (D) Percentages of mG-peak, mN-peak, mTB-peak, and Lv-peak genes in two gene populations; genes under control of *Meox*, *Tcf15-r*, and/or *Tbx15-18/22*, and genes under control of none of them.

The *Meox* MO used in the above experiment was designed to block translation. To confirm its specificity, we designed another MO, which was expected to block splicing. This splicing-block MO indeed blocked splicing, although its efficiency was not very high (Figure S5A). Next, we examined the expression level of *Ckmt*, because it was the most prominently downregulated gene in the RNA-seq experiment using embryos injected with the *Meox* translation-block MO (Figure 4B). RT-qPCR experiments showed that this gene was downregulated in embryos injected with the splicing-block MO similarly to embryos injected with the translation-block MO. Likewise, a splicing-block MO against *Tcf15-r* indeed blocked splicing (Figure S5C), and *Acta.h*, which encodes muscle actin, was downregulated in embryos injected with the translation-block MO and in embryos injected with the splicing MO against *Tcf15-r* (Figure S5D). For this experiment, we used primers designed for an intron of *Acta.h* and measured expression levels of its nascent transcript, because the ascidian genome contains several muscle actin genes and their exons are highly similar. These observations indicated that all these MOs specifically knocked down *Meox* and *Tcf15-r*.

According to temporal expression profiles based on the RNA-sequencing data (Brozovic et al., 2018), five of six downstream genes of *Meox* were Lv-peak genes and the remaining one was an mTB-peak gene (Figure S6). Similarly, all nine downstream genes of *Tcf15-r* were Lv-peak genes (Figure S6). Among the 42 genes regulated by *Tbx15/18/22*, 33 genes were Lv-peak genes, and 9 genes were mTB-peak genes (Figure S6). On the other hand, the remaining 54 genes, which were not under control of *Meox*, *Tcf15-r*, or *Tbx15/18/22*, showed quite a different pattern from those under control of *Meox*, *Tcf15-r*, and/or *Tbx15/18/22* (Figure 4D; Fisher’s exact test p-value, 5.5×10^-7^). This observation indicates that *Meox*, *Tcf15-r*, and *Tbx15/18/22* individually or combinatorically maintain expression levels in late embryos and larvae.

### Snai regulates muscle-specific genes in earlier stages

Next, we examined *Snai* morphants at the middle tailbud stage by RNA-sequencing. Because the *Snai* MO was used in previous studies (Imai et al., 2006; Imai et al., 2009; Tokuoka et al., 2018), we did not examine specificity of this MO in the present study. We found that 43 genes were significantly down-regulated (adjusted p-values, <0.01) (Figure 5A; Table S2). The temporal expression profile of *Snai* shown in Figure 3 suggests that *Snai* may be involved in regulation of genes at earlier stages than *Meox*, *Tcf15-r*, and *Tbx15/18/22*. Indeed, *Tcf15-r* and *Meox*, which are expressed most abundantly at the middle tailbud stage (see Figure 3), were downregulated in *Snai* morphants (Figure 5A). To further test this hypothesis, we compared expression profiles of genes under control of *Snai* with those of genes that are not. Temporal expression profiles of genes regulated by *Snai*, but not by *Meox*, *Tcf15-r*, or *Tbx15/18/22* were markedly different from those of genes regulated by *Meox*, *Tcf15-r*, or *Tbx15/18/22* but not by *Snai* (Figure 5C). Their profiles also differed from those of genes not under control of *Snai*, *Meox*, *Tcf15-r*, or *Tbx15/18/22* (Figure 5C; Figure S7).

**Figure 5.**
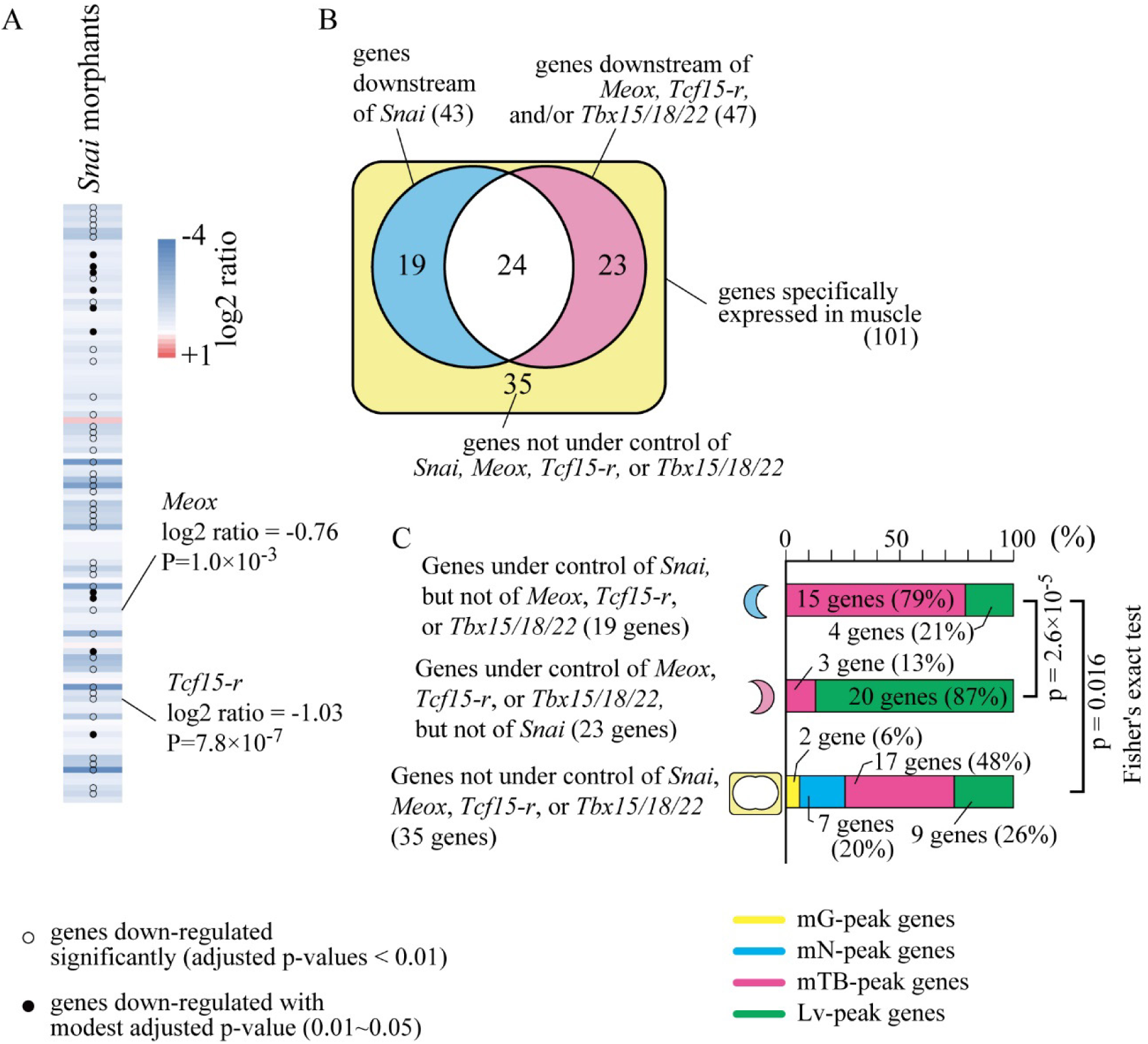
*Snai* regulates a subset of muscle-specific genes, most of which are expressed most abundantly in middle tailbud embryos. (A) A heatmap of changes in expression of the 101 muscle-specific genes between control and *Snai* morphant embryos. RNA-sequencing results were analyzed with Deseq2 (Love et al., 2014). Genes, expression of which was significantly downregulated, are shown with dots. (B) A Venn diagram that shows overlaps between genes under control of *Snai* and genes under control of *Meox*, *Tcf15-r*, and/or *Tbx15/18/22*. (C) Percentages of genes with peak expression at four developmental stages in three gene populations: genes under control of *Snai* but not of *Meox*, *Tcf15-r*, or *Tbx15-18/22*, genes under control of *Meox*, *Tcf15-r*, and/or *Tbx15-18/22* but not of *Snai*, and genes under control of none of them.

### Muscle-specific transcription factors combinatorically regulate temporal expression patterns of downstream genes

The above analysis indicated that temporal expression profiles of muscle-specific genes are determined by combinations of upstream regulatory factors and encoded in cis-regulatory regions of downstream genes. Because the above analysis indicated that *Snai* promotes expression around the tailbud stage, we next tried to confirm that Snai binding sites can promote expression around the tailbud stage using reporter constructs. For this purpose, we first examined the upstream regulatory region of *Meox* and *Tcf15-r*. The aforementioned RNA-sequencing analyses showed that *Meox* and *Tcf15-r* are under control of *Snai* (Figure 5A; Table S2), and we identified two putative Snai binding sites for each of these genes using JASPAR scan (Sandelin et al., 2004) and the *Ciona* Snai position weight matrix (Nitta et al., 2019) (Figure S8). Upstream regions of *Meox* and *Tcf15-r* containing these putative Snai binding sites promoted reporter expression in muscle cells of middle tailbud embryos (Figure 6A, B). Using gel-shift assays, we confirmed that these putative Snai binding sites indeed bound Snai (Figure 6C).

**Figure 6.**
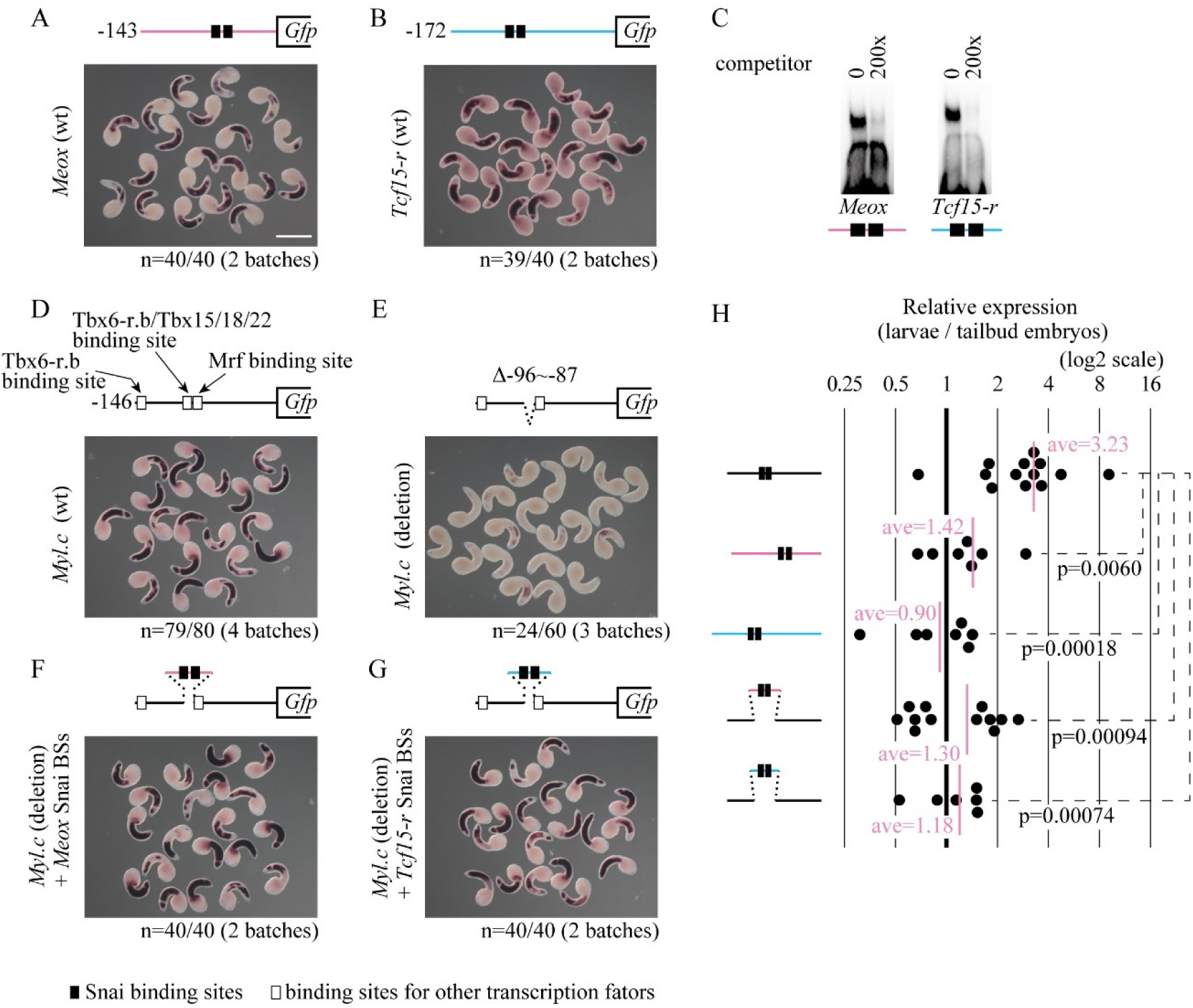
*Snai* binding sites promote expression around the middle tailbud stage. (A, B, D–G) Expression of a reporter gene (*GFP*) was examined by *in situ* hybridization at the middle tailbud stage. (A, B) 143-bp and 172-bp upstream regions of *Meox* (A) and *Tcf15-r* (B), which contain two Snai binding sites, are sufficient to promote expression of the reporter in muscle cells. (C) Gel shift assays show that upstream regions of *Meox* and *Tcf15-r* can bind Snai *in vitro*. Shifted bands disappear by addition of specific competitors, indicating specific binding of Snai to these regions. (D) The 146-bp upstream region of *Myl.c*, which contains a Tbx6-r.b binding site, a shared binding site for Tbx6-r.b and Tbx15/18/22, and E-box (Mrf binding site), is sufficient to promote the reporter in muscle cells, as reported (Yu et al., 2019). (E) Deletion of the shared binding site for Tbx6-r.b and Tbx15/18/22, which is important for expression in tailbud embryos and larvae (Yu et al., 2021; Yu et al., 2019), almost abolished expression in muscle cells at the middle tailbud stage, although a small number of embryos expressed the reporter very weakly. (F, G) Insertions of 51-bp and 45-bp upstream regions of *Meox* and *Tcf15-r* containing two *Snai* binding sites into the *Myl.c* deletion construct recovered expression of the reporter in muscle cells. (H) Expression levels of the reporters shown in (A), (C), (E), (F), and (G) were compared between the middle tailbud and larval stages, and relative expression levels are shown. Each dot represents an independent experiment, and bars show averages. Differences were tested by Wilcoxon’s rank sum test, and p-values are shown. Scale bar, 200 μm.

A previous study demonstrated that *Myl.c*, whose expression peak was at the larval stage (Table S3), has two cis-regulatory modules in its 146-bp upstream region that can promote reporter expression at the tailbud stage (Figure 6D; Figure S8) (Yu et al., 2021; Yu et al., 2019). Deletion of the shared binding site for Tbx6-r.b and Tbx15/18/22 greatly reduced the activity of this regulatory region, because only 24 of 60 experimental embryos expressed the reporter very weakly (Figure 6E). We next inserted 51-bp and 45-bp upstream regions of *Meox* and *Tcf15-r* that contained Snai binding sites instead of the shared binding site for Tbx6-r.b and Tbx15/18/22. These reporter constructs were expressed in muscle cells of middle tailbud embryos (Figure 6F, G). Therefore, the inserted *Snai* sites promoted expression in middle tailbud stage.

Finally, we compared expression levels of the reporter genes in middle tailbud embryos with those in larvae using RT-qPCR (Figure 6H). The expression level of the reporter with the wild-type upstream region of *Myl.c* increased greatly from the middle tailbud stage to the larval stage, while those of the reporters with wild-type upstream regions of *Meox* and *Tcf15-r* were rarely changed. Expression levels of the chimera construct, which were tested in the above experiments (Figure 6F and G), were rarely changed between the middle tailbud and larval stages. That is, replacements of shared binding sites for *Tbx6-r.b* and *Tbx15/18/22* with *Snai* binding sites of the upstream regions of *Meox* and *Tcf15-r* shifted the expression peak earlier, as expected.

## Discussion

### The regulatory circuit for differentiation of muscle cells of ascidian larvae

Figure 7 summarizes the genetic pathway for differentiation of B-line muscle cells of ascidian larvae. The muscle determinant, Zic-r.a or Macho-1, is primarily responsible for initiation of this pathway (Nishida and Sawada, 2001; Satou et al., 2002). On the other hand, another Zic-related transcription factor gene, *Zic-r.b*, is activated by a pathway under control of β-catenin and Gata.a, but independently of Zic-r.a (Imai et al., 2016; Imai et al., 2002; Tokuoka et al., 2021). Similarly, *Snai* is also activated in a subset of B-line muscle cell progenitors through maternally expressed, constitutively active Raf kinase, which also acts independently of Zic-r.a, while *Snai* is activated in the remaining B-line muscle cell progenitors by *Tbx6-r.b* under control of Zic-r.a (Fujiwara et al., 1998; Tokuoka et al., 2018). *Tbx6-r.b* activates *Mrf*, and these two genes constitute a feedback loop (Yu et al., 2019). All 101 muscle-specific genes examined in the present study are under control of this circuit.

**Figure 7.**
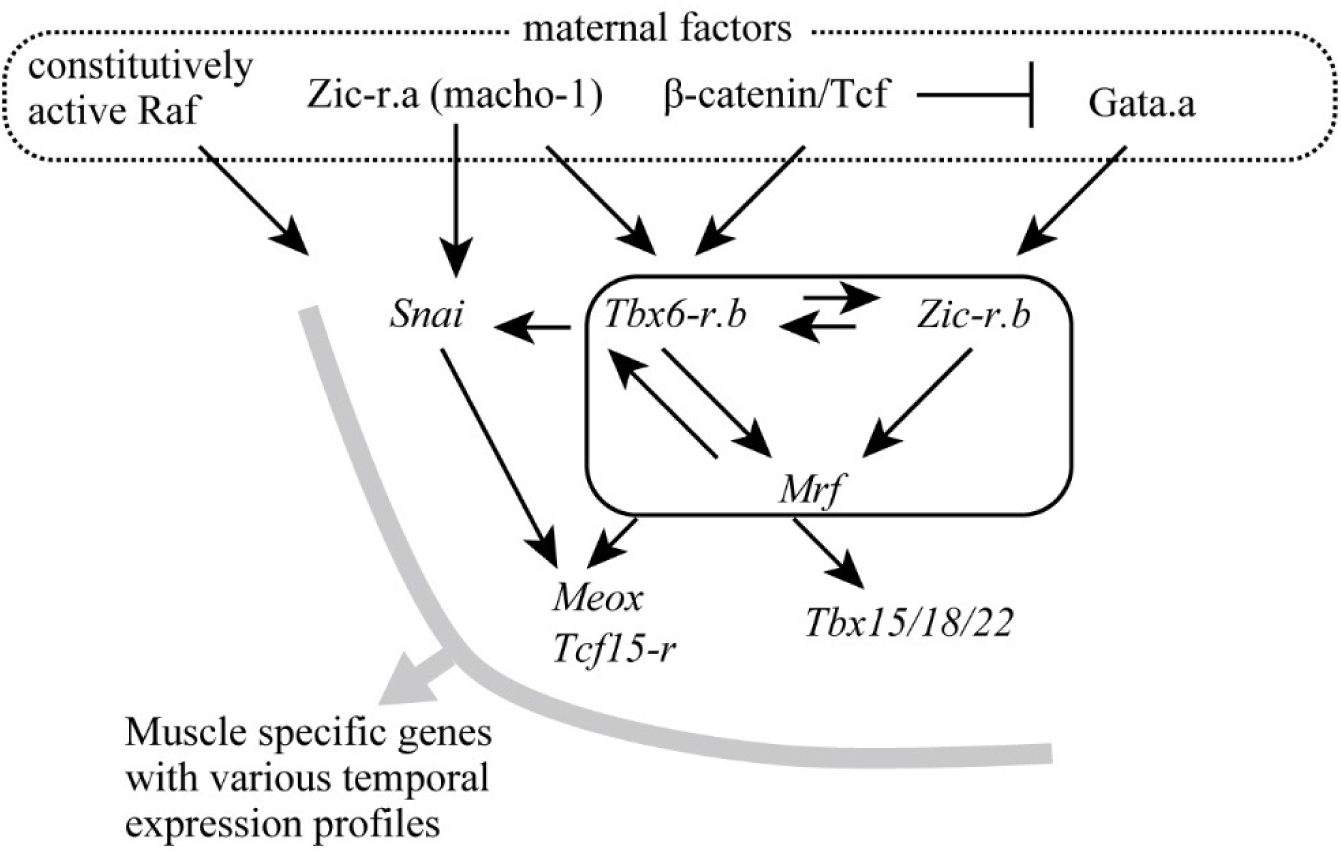
The gene regulatory circuit from maternal factors to muscle structural genes in ascidian larval muscle development. The present study indicates that all muscle-specific genes are under control of this gene circuit. Muscle-specific genes have various temporal expression profiles that are likely important for muscle development. These temporal expression profiles are created by this gene regulatory circuit.

*Tbx15/18/22*, which is activated under control of the regulatory loop of *Tbx6-r.b* and *Mrf*, maintains expression of *Myl.c* by the larval stage (Yu et al., 2021). Here, we provided evidence that *Meox* and *Tcf15-r* also helps to maintain expression of a subset of muscle-specific genes, which partly overlaps with target genes of *Tbx15/18/22*.

In addition, many muscle-specific genes are abundantly expressed in early embryos and rarely expressed in late embryos or larvae (see Figure 2). *Meox* and *Tcf15-r* are such genes, and controlled by *Snai*, which is responsible for expression at the tailbud stage. Thus, in the muscle lineage of ascidian embryos, a combination of Tbx6-r.b, Mrf, Snai, Meox, Tcf15-r, and Tbx15/18/22 coordinates temporal expression patterns of muscle-specific genes.

Although *Snail* encodes a well-known transcriptional repressor (Chopra and Levine, 2009), which was originally identified in *Drosophila* (Simpson, 1983), this protein can also activate targets in a context dependent manner (Rembold et al., 2014). We showed that Snail participates in activating its targets in muscle cells of ascidian embryos, whereas a previous study suggests that Snail represses *Brachyury*, an essential transcription factor for specification of the notochord (Fujiwara et al., 1998).

In addition, Mrf protein levels are controlled by cereblon, a substrate receptor of an E3 ubiquitin ligase, and this helps to control temporal expression profiles of muscle-specific genes (Long et al., 2023). It also contributes to temporal regulation of muscle development. Importantly, *cereblon* is also under control of Mrf (Long et al., 2023).

### Temporal regulation of muscle-specific genes

We have showed that various temporal expression patterns are reproducible and can be designed. Reporters with only Tbx6-r.b-binding sites are expressed in early embryos, and initiation timing is controllable via the number of Tbx6-r.b-binding sites (Yu et al., 2019). Reporters with a Tbx6-r.b binding site and a Snai-binding site are expressed in early embryos and tailbud embryos. Reporters with a Tbx15/18/22 site are expressed even in larvae (Yu et al., 2021).

Importantly, these expression profiles reproduce those of endogenous genes. Indeed, genes encoding myofiber proteins tend to be expressed later and other genes tend to be expressed earlier, which suggests that temporal regulatory mechanisms are essential for differentiating functional muscle cells.

This temporal regulatory mechanism is similar to that in the notochord lineage, in which *Brachyury* is specifically expressed (Corbo et al., 1997; Yasuo and Satoh, 1993). Genes with multiple Brachyury binding sites that can act synergistically are expressed earlier. Genes with a single Brachyury binding site are expressed next, and then genes that are indirectly controlled by Brachyury are expressed (Katikala et al., 2013). Temporal regulation of gene expression by combinations of transcription factors, and numbers and probably qualities of their binding sites is likely a common mechanism in various tissues of ascidian larvae.

### Secondary muscle cell lineages

In addition to B-line muscle cells, ascidian larvae have four pairs of muscle cells that are not differentiated autonomously, and these cells are called A-line and b-line muscle cells, because they are derived from A4.1 and b4.2 blastomeres of 8-cell embryos, respectively. In A-line cells, *Tbx6-r.b* and *Mrf* are expressed under control of FGF, Delta/Notch, and Nodal signaling pathways (Hudson et al., 2007). In the b-line, *Tbx6-r.b* is also expressed soon after fate restriction at the early neurula stage (Ishida and Satou, 2024). Although it has not been determined precisely when *Mrf* begins to be expressed in this lineage, *Mrf* is clearly expressed in b-line muscle cells of tailbud embryos (Meedel et al., 1997). These observations indicate that *Tbx6-r.b* and *Mrf* are also responsible for differentiation of A-line and b-line muscle cells, and our RNA-sequencing analysis using morphants of *Tbx6-r.b* and *Mrf* supports this hypothesis.

This means that all three lineages of ascidian larval muscle cells share the gene circuit that begins with the regulatory loop of *Tbx6-r.b* and *Mrf*. Although it is currently unknown which lineage represents an ancestral pathway, it is likely that the regulatory loop of *Tbx6-r.b* and *Mrf* and its downstream circuit has been co-opted from an ancestral lineage to other two lineages, as in the case of dorsal and ventral epidermal sensory neurons of ascidian larvae (Waki et al., 2015).

### Tbx6-r.b and Mrf constitute the master regulatory loop for muscle differentiation

We identified 101 muscle-specific genes using single-cell transcriptome data, and almost all known genes expressed in muscle cells are included in this list. Therefore, it is likely that almost all genes specifically expressed in muscle cells were examined in the present study. Even if a small number of genes failed to be identified for technical reasons, it is likely that all muscle-specific genes including such missing genes are under control of the regulatory loop of *Tbx6-r.b* and *Mrf*.

In addition to these muscle-specific genes, there are genes that are expressed in multiple cell types including muscle cells (Satou et al., 2001). Although it is not easy to analyze these genes exhaustively, they often have multiple cis-regulatory modules that direct tissue-specific gene expression. Therefore, it is likely that cis-regulatory modules that induce muscle-specific expression are similarly under control of *Tbx6-r.b* and *Mrf*.

As we showed previously (Yu et al., 2019), *Myl.c* is directly controlled by *Tbx6-r.b* and *Mrf*. Similar cis-regulatory sequences, which are expected to bind Tbx6-r.b and Mrf, are found in many other muscle-specific genes (Satou and Satoh, 1996; Yuasa et al., 2002). Therefore, these studies altogether indicate that that the regulatory loop of *Tbx6-r.b* and *Mrf,* but not *Mrf* alone, acts as the master regulator for muscle differentiation in ascidian embryos.

This mechanism contrasts with that in vertebrate embryos, in which myogenic factors serve the master regulatory function. Interestingly, four myogenic factor genes in vertebrate genomes regulate one another (Tapscott, 2005). For example, Myod and myogenin can activate each other (Brennan et al., 1990; Thayer et al., 1989). Such cross-regulatory circuits may function in a manner equivalent to that of the regulatory loop we found in ascidian embryos. If so, acquisition of paralogous genes in the vertebrate lineage may have promoted a change in the structure of the gene regulatory network controlling muscle development, and these paralogs acquired a genuine master regulatory function in the vertebrate lineage.

## Materials and Methods

### Animals and cDNAs

*Ciona robusta* (or *C. intestinalis*, type A) adults were obtained from the National Bio-Resource Project for *Ciona intestinalis*. cDNA clones were obtained from our EST clone collection (Satou et al., 2005). Identifiers for genes examined in the present study are listed in Table S6.

### Whole-mount in situ hybridization and RT-qPCR to detect expression of endogenous genes

*In situ* hybridization was performed as described previously (Ikuta and Saiga, 2007; Satou et al., 1995). For experiments shown in Figure 3, we extracted RNA from 50 embryos using an RNeasy Micro kit (Qiagen, #74004). Before extraction, we mixed synthetic *Gfp* mRNA as a spike-in control, and used it to normalize differences in extraction efficiency among samples, because all genes expressed at a constant level from eggs to larvae have been identified. Normalized expression values per embryo were compared with those at the late neurula stage. We similarly performed experiments shown in Figures S2, S3, and S5, but we did not use spike-in controls for these experiments. We instead used *Raf* as an internal control for relative quantification. *Cdk5* and *Slk/Stk10* (Tokuoka et al., 2022) were used as additional controls. SYBR green chemistry was used for qPCR.

Primers for the experiment are shown in Table S7. There are 10 genes encoding muscle actin in the genome (*Acta.a* to *Acta.j*). Primers for the experiment shown in Figure S2 were expected to detect expression of all actin genes, whereas primers for the experiment shown in Figure S5 were designed to specifically amplify an intronic region of *Acta.h* to specifically detect nascent *Acta.h* transcripts. We also used primers to amplify an intronic region of *Myl.c* to detect nascent *Myl.c* transcripts.

### Gene knockdown assays and reporter assays

MOs (Gene Tools, LLC) used in the present study are listed in Table S8. The *Mrf* MO and a *Tbx6-r.b* MO (5’-TTACAATTTCCTCTCTCTTTCGATT-3’) have been used in previous studies; therefore, we did not perform experiments to further confirm their specificity in the present study. Specificity of a newly designed *Tbx6-r.b* MO (5’-TCATATTCGCCATAGTCTTGTCTGG-3’), and MOs against *Meox* and *Tcf15-r* were tested as described in Results (Figure S2 and Figure S5). We also used a MO against *lacZ* of *Escherichia coli* as a control. MOs were introduced by microinjection under a microscope. All knockdown phenotypes were confirmed in at least two independent series of experiments.

Genomic positions of the upstream regions of *Meox, Tcf15-r*, and *Myl.c* are 6,019,222– 6,019,487 on chromosome 8, 250,326–250,558 on chromosome 11, and 5,405,563–5,405,768 on chromosome 8 of the HT assembly (Satou et al., 2019), respectively. The reporter constructs contain a genomic region of the first and second exons and the first intron of *Foxtun2* (Chromosome 14: 6279296–6279458) before GFP, to avoid amplification from introduced DNAs in qPCR (Figure S9; see below). Reporter constructs were introduced into fertilized eggs by electroporation. Expression of the reporter was examined by *in situ* hybridization and by RT-qPCR.

For RT-qPCR to detect reporter expression, we used a Cell-to-Ct kit (Thermo-Fisher Scientific, #A35377) to obtain cDNA from 20 to 30 embryos. Because we did not have good controls expressed at the same level between the tailbud and larval stages, we did not use internal controls, but directly compared expression levels between the same number of tailbud embryos and larvae from the same batch. We used the Taqman method for qPCR. Primers and a probe were designed to amplify cDNA derived from spliced mRNA (Forward primer: 5’-CAGTTCGTTTAAAG GG-3’, Reverse primer: 5’-GCCGGACACGCTGAACTT-3’, Probe: 5’-FAM-CTGGTCGAGCTGGACGGCGAC-TAMRA-3’). To ensure preferential amplification of cDNA from spliced mRNA, we changed the reaction conditions as follows: primer concentrations, 300 nM each; probe concentration, 250 nM; extension duration, 10 seconds.

### Gel-shift assay

Recombinant Snai protein was produced as a fusion protein of the Snai DNA-binding domain (amino acid positions: 436–581) and glutathione S-transferase in *Escherichia coli* DH5α strain. The protein was purified under native conditions using MagneGST Glutathione Particles (Promega, #V861A). After annealing two complementary oligonucleotides (5’-AAAATACGAGGTCAGTCGTCACCTTTGCTTGCCCAGTTGTTTACTTCGTTTAAA -3’ and 5’-AAATTTAAACGAAGTAAACAACTGGGCAAGCAAAGGTGACGACTGACCTCGTAT -3’ for the *Meox* upstream region; 5’-AAACGTGTAGTTCAAGACAGCTGACTCACGTGATATCAGCATGAATGG -3’ and 5’-AAACCATTCATGCTGATATCACGTGAGTCAGCTGTCTTGAACTACACG -3’ for the *Tcf15* upstream region), both protruding ends of the double-stranded oligonucleotides were filled with biotin-11-dUTP, and these biotin-labelled oligonucleotides were used as probes. Proteins and biotin-labeled probes were mixed in 25 mM Hepes (pH 7.5), 50 mM KCl, 1 mM DTT, 25 ng/μL poly(dIdC), 2.5% glycerol, 0.05% NP40, and 100 ng/μL BSA with or without competitor double-stranded DNAs (200-fold molar excess). Protein amounts were determined empirically. Protein-DNA complexes were detected using a Chemiluminescent Nucleic Acid Detection Module Kit (Thermo Fisher Scientific, #89880).

### RNA-sequencing

For RNA-sequencing, we prepared embryos injected with the *lacZ* MO (0.5 mM), *Tbx6-r.b* MO (0.5 mM), *Mrf* MO (0.5 mM each), *Snai* MO (0.5 mM), *Meox* translation-block MO (0.5 mM), or *Tcf15-r* translation-block MO (0.5 mM). We also prepared embryos injected with *Tbx6-r.b* and *Mrf* simultaneously (0.5mM each), and their controls injected with a double dose of the *lacZ* MO (1 mM). Total RNA was extracted using a Dynabeads mRNA DIRECT Micro Kit (Thermo Fisher Scientific, #61011). For *Mrf* morphants, we made libraries with an Ion Total RNA-Seq kit version 2 (Thermo Fischer Scientific, #4475936), and used Ion PGM/S5XL instruments (DRA accession numbers: DRR585068, DRR585069). These embryos were collected when we collected *Tbx15/18/22* morphants and control embryos in our previous study (Yu et al., 2021). Therefore, controls for *Mrf* morphants were those that we previously used to analyze *Tbx15/18/22* morphants [DRA accession numbers: DRR315181 and DRR315182 (Yu et al., 2021)]. For the remaining samples, libraries were made with an NEBNext Ultra II Directional RNA Library Prep kit (New England Biolabs, #7765). Obtained reads were deposited in the DRA/SRA database under accession numbers: DRR585933–DRR585950. The experiment included two biological replicates. Sequencing reads were mapped to the HT-version of the genome (Satou et al., 2019) and the latest gene model set (Satou et al., 2022) using bowtie2 (Langmead and Salzberg, 2012) with default parameters, and DESeq2 (Love et al., 2014) was used to identify differentially expressed genes with default parameters. RNA-sequencing data of wild type embryos (Brozovic et al., 2018) were mapped in the same way. Single-cell transcriptome data (Cao et al., 2019) were mapped to the same set of the genome and gene models using Cell Ranger (10xGenomics) with default parameters.

*Ciona* muscle-specific proteins were mapped to the human proteome (Uniprot release 2021_01, file name: UP000005640_9606.fasta) (UniProt-Consortium, 2019) using BLASTP with default parameters (Altschul et al., 1990). Best hit proteins with E-values <1×10^-5^ were regarded as homologs. GO annotations were downloaded from the Gene Ontology web site (file name: goa_human.gaf, downloaded at June 10^th^, 2024) (Ashburner et al., 2000).

## Acknowledgements

We thank Chikako Imaizumi, Manabu Yoshida and other members working under the National Bio-Resource project (MEXT, Japan) in my group and the University of Tokyo for providing experimental animals.

## Competing interests

The authors declare no competing or financial interests.

## Funding

This research was supported by grants from the Japan Society for the Promotion of Science (21H02486, 24K02032, 24K21274) to YS and (21K06017) to IO, and from Takeda Science Foundation to YS.

## Data availability

RNA-sequencing data produced in the present study are available in the DRA/SRA database under accession numbers DRR585068, DRR585069, and DRR585933–DRR585950.

---

**Figure S1.**
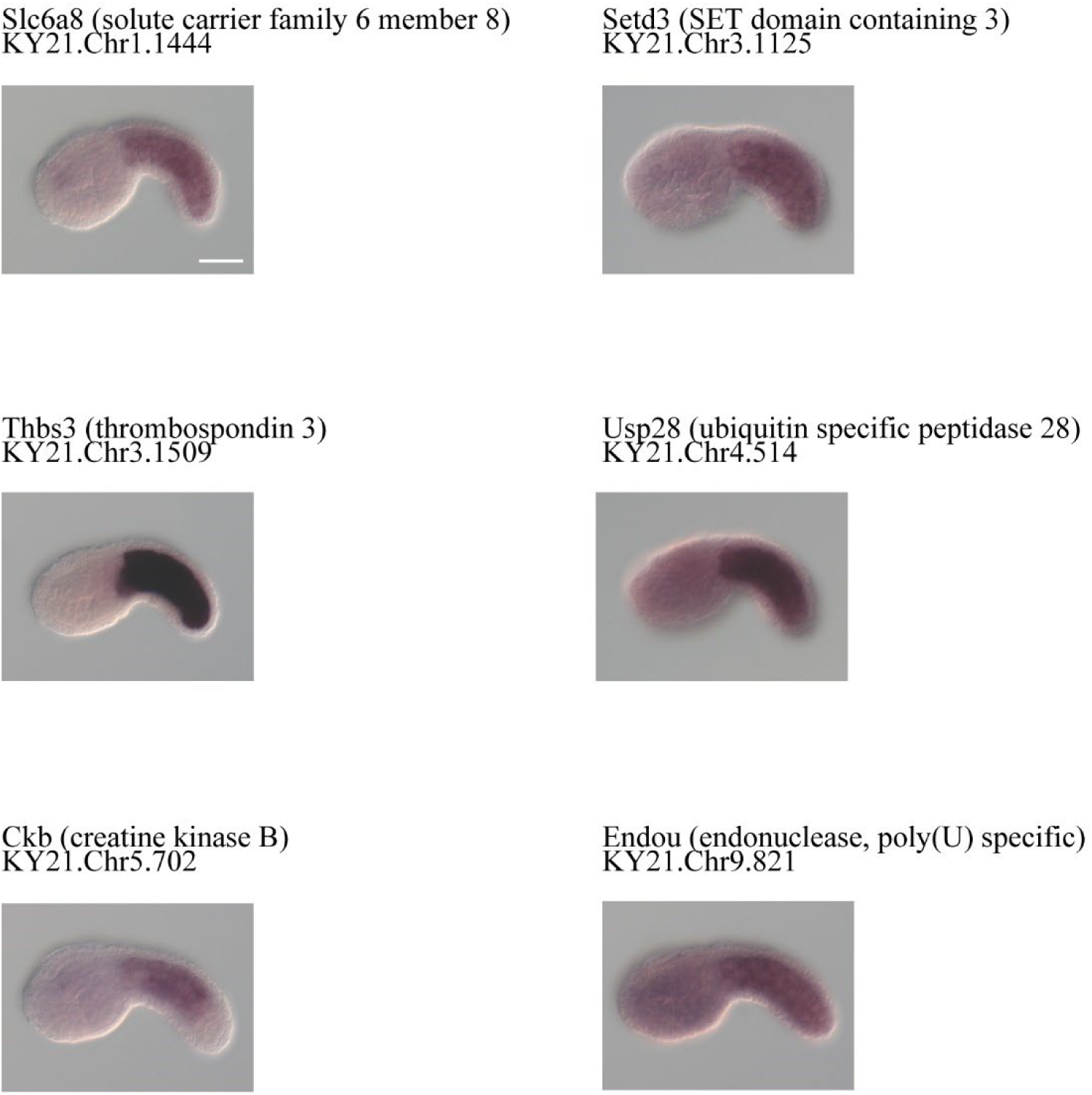
Expression of genes randomly chosen among genes that were identified as muscle-specific genes by an analysis of single-cell transcriptome data, but had not been examined by *in situ* hybridization. Gene product names and gene identifiers are shown above the photographs. Scale bar, 50 μm.

**Figure S2.**
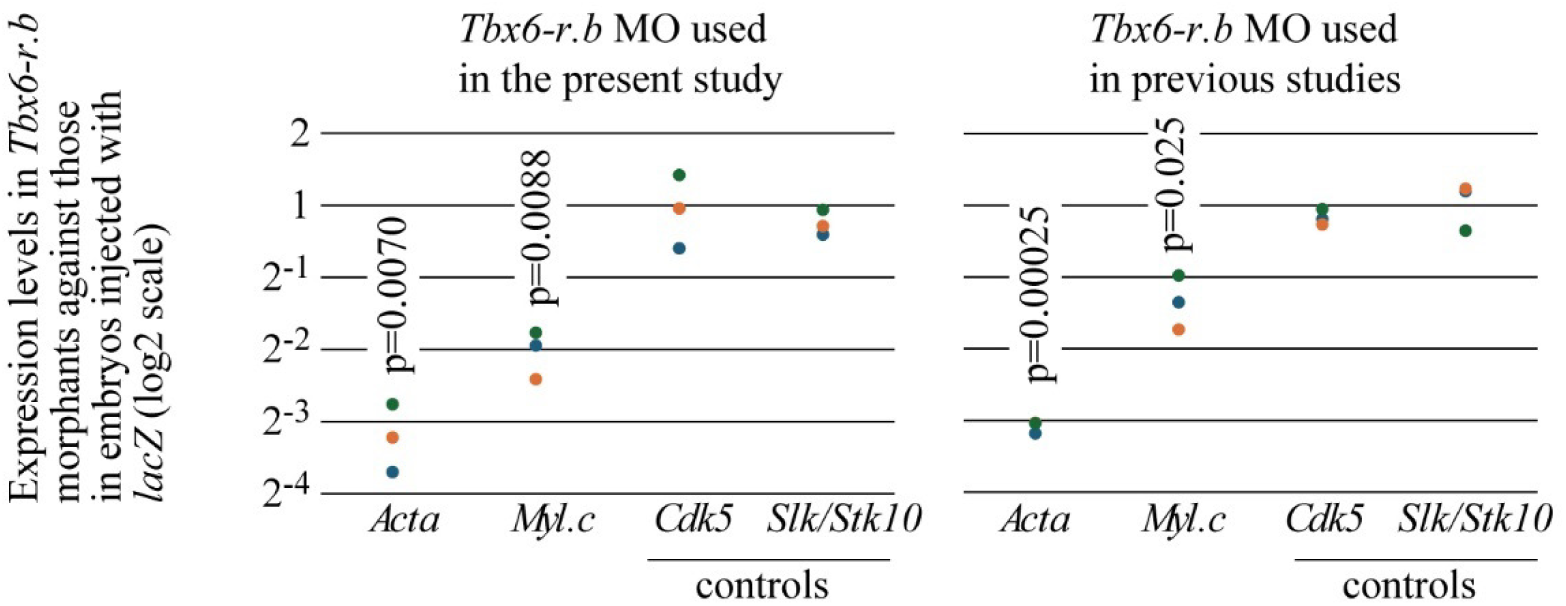
Confirmation of specificity of a newly produced MO against *Tbx6-r.b*. Expression levels of muscle actin and *Myl.c* genes were measured by RT-qPCR. *Raf* was used as an internal reference for normalization between samples, and *Cdk5* and *Slk/Stk10* were used as negative controls. The y-axis indicates relative expression levels of embryos injected with the new *Tbx6-r.b* MO or the previously used *Tbx6-r.b* MO against control embryos injected with *lacZ* MO. Results from three batches of embryos are shown in two colors. P-values were calculated by paired t-tests using Ct values normalized with *Raf* expression levels. These tests indicate that different expression levels of genes encoding muscle actin and myosin light chain (*Myl.c*) between *Tbx6-r.b* morphants and embryos injected with the *lacZ* MO were significant in both cases. Muscle-actin gene primers were designed in exons and expected to produce amplicons for all muscle actin genes, whereas primers for *Myl.c* were designed to amplify an intronic region for detection of nascent transcripts.

**Figure S3.**
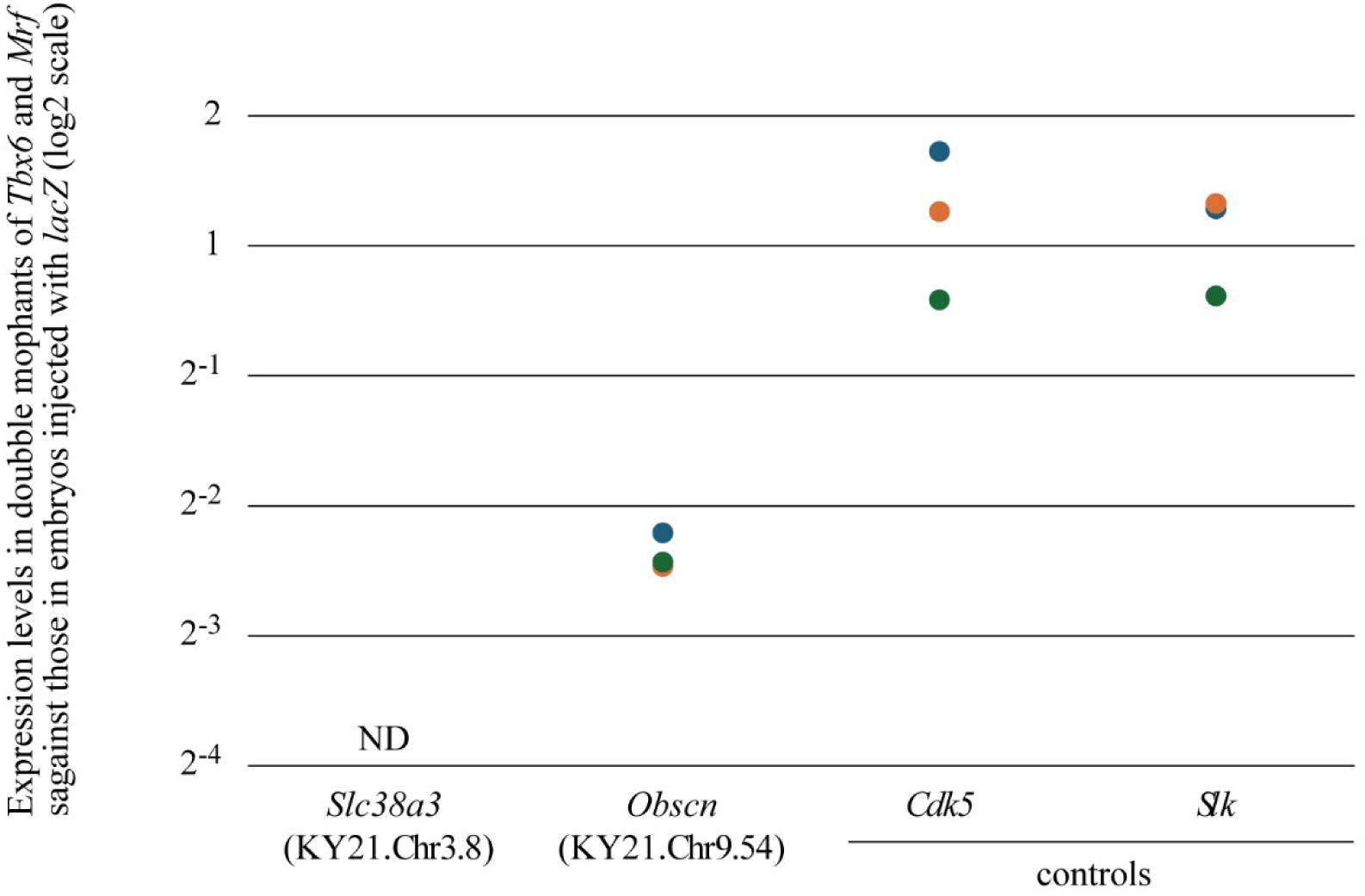
Downregulation of *Slc38a3* and *Obscn* in double morphants of *Tbx6-r.b* and *Mrf.* Expression levels of *Slc38a3* (KY21.Chr3.8) and *Obscn* (KY21.Chr9.54) were measured by RT-qPCR, because down-regulation of these two genes in double morphants of *Tbx6-r.b* and *Mrf* was indicated only moderately in the RNA-sequencing experiment. *Raf* was used as an internal reference for normalization between samples, and *Cdk5* and *Slk/Stk10* were used as negative controls. The y-axis indicates relative expression levels of double morphants of *Tbx6-r.b* and *Mrf* against control embryos injected with *lacZ* MO. Results from three batches of embryos are shown in three colors. Differences were examined by paired t-tests using Ct values normalized with *Raf* expression levels. Expression levels of *Obscn* were significantly different between double morphant embryos of *Tbx6-r.b* and *Mrf* and embryos injected with the *lacZ* MO (p-value, 0.0012). Because *Slc38a3* expression was rarely detected in double morphant embryos of *Tbx6-r.b* and *Mrf*, its p-value cannot be calculated.

**Figure S4.**
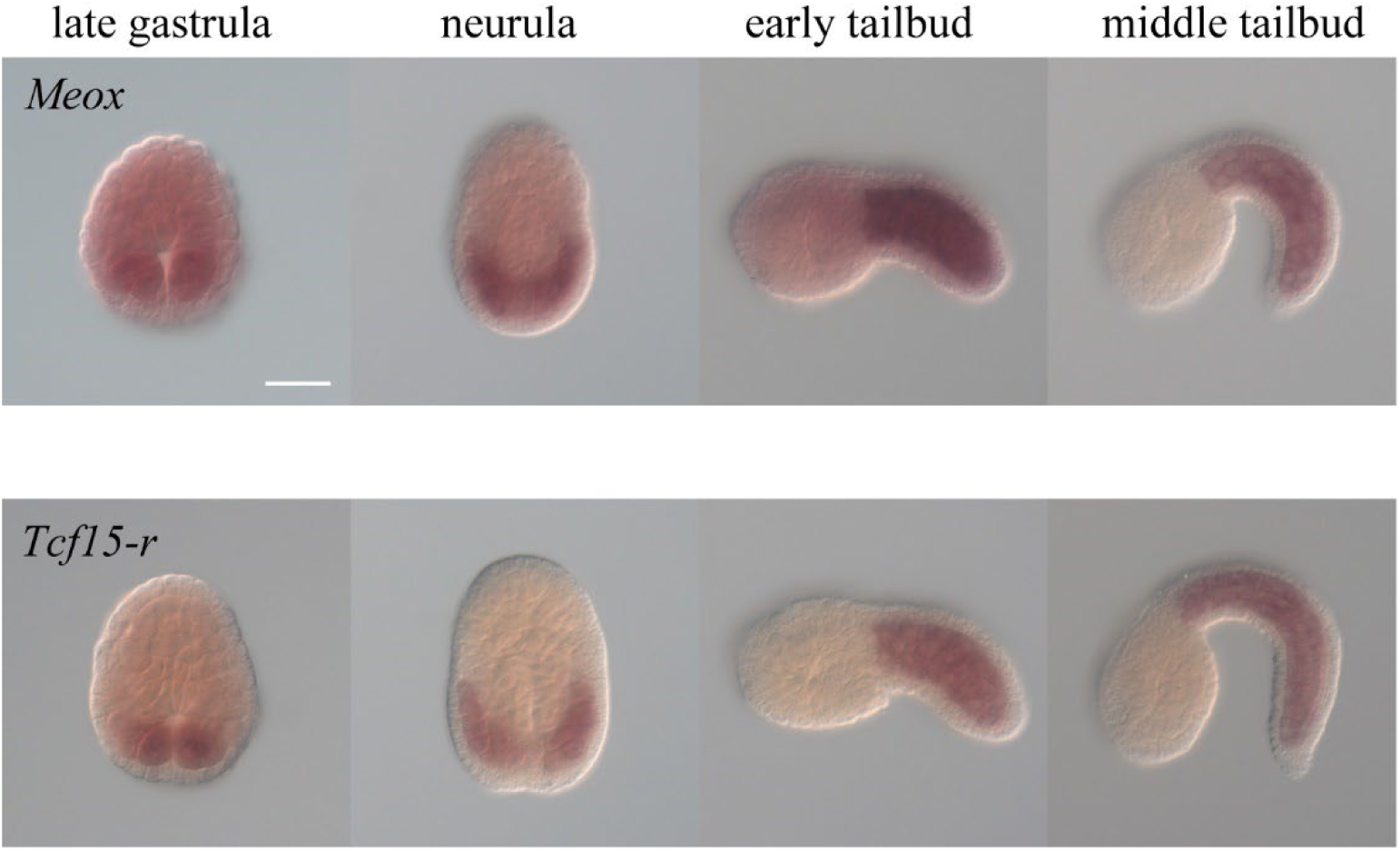
Spatial and temporal expression patterns of *Meox* and *Tcf15-r* revealed by *in situ* hybridization. Both *Meox* and *Tcf15-r* initiate expression at the late gastrula stage in the muscle lineage. Scale bar, 50 μm.

**Figure S5.**
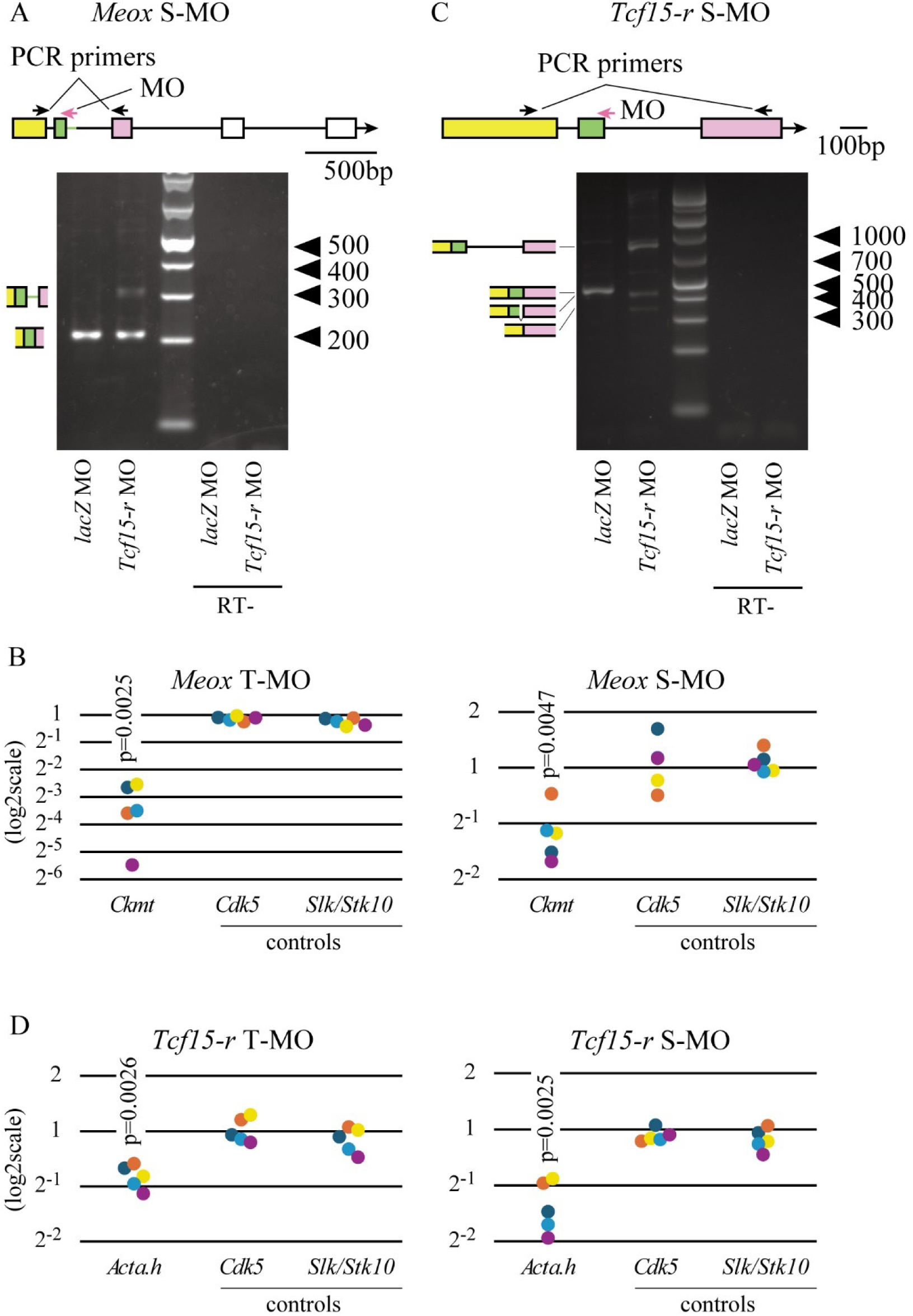
Confirmation of specificity of MOs against *Meox* and *Tcf15-r*. (A–D) We designed MOs that were expected to block translation (T-MOs) and MOs that were expected to block proper splicing (S-MOs). (A, C) To confirm that S-MOs successfully block splicing, RNA extracted from experimental embryos was reverse-transcribed, and obtained cDNA pools were used as templates for subsequent PCR. Positions of PCR primers and S-MOs are indicated at the top. While a single band was amplified from control embryos injected with *lacZ* MO in both cases, multiple bands were amplified from embryos injected with (A) *Meox* S-MO or (C) *Tcf15-r* S-MO. These bands were extracted from gels, and their nucleotide sequences were determined. Deduced structures are depicted on the left. No bands were amplified from controls, in which water was added instead of reverse transcriptase (RT-). (B, D) We measured expression levels of (B) *Ckmt* (KY21.Chr4.755) in embryos injected with *Meox* MOs, and (D) *Acta.h* (KY21.Chr1.2020) for *Tcf15-r* MOs. To discriminate *Acta.h* from other muscle actin genes, we used primers designed to amplify an intronic region of *Acta.h*; therefore, we measured expression levels of nascent transcripts of *Acta.h*. Expression levels are shown as relative values against expression levels in control embryos injected with *lacZ* MO. Results from five different batches of embryos are shown in five colors. P-values shown in the graphs were calculated by paired t-tests using Ct values normalized with *Raf* expression levels. These tests indicate that differences in expression levels of *Ckmt* and *Acta.h* were significant.

**Figure S6.**
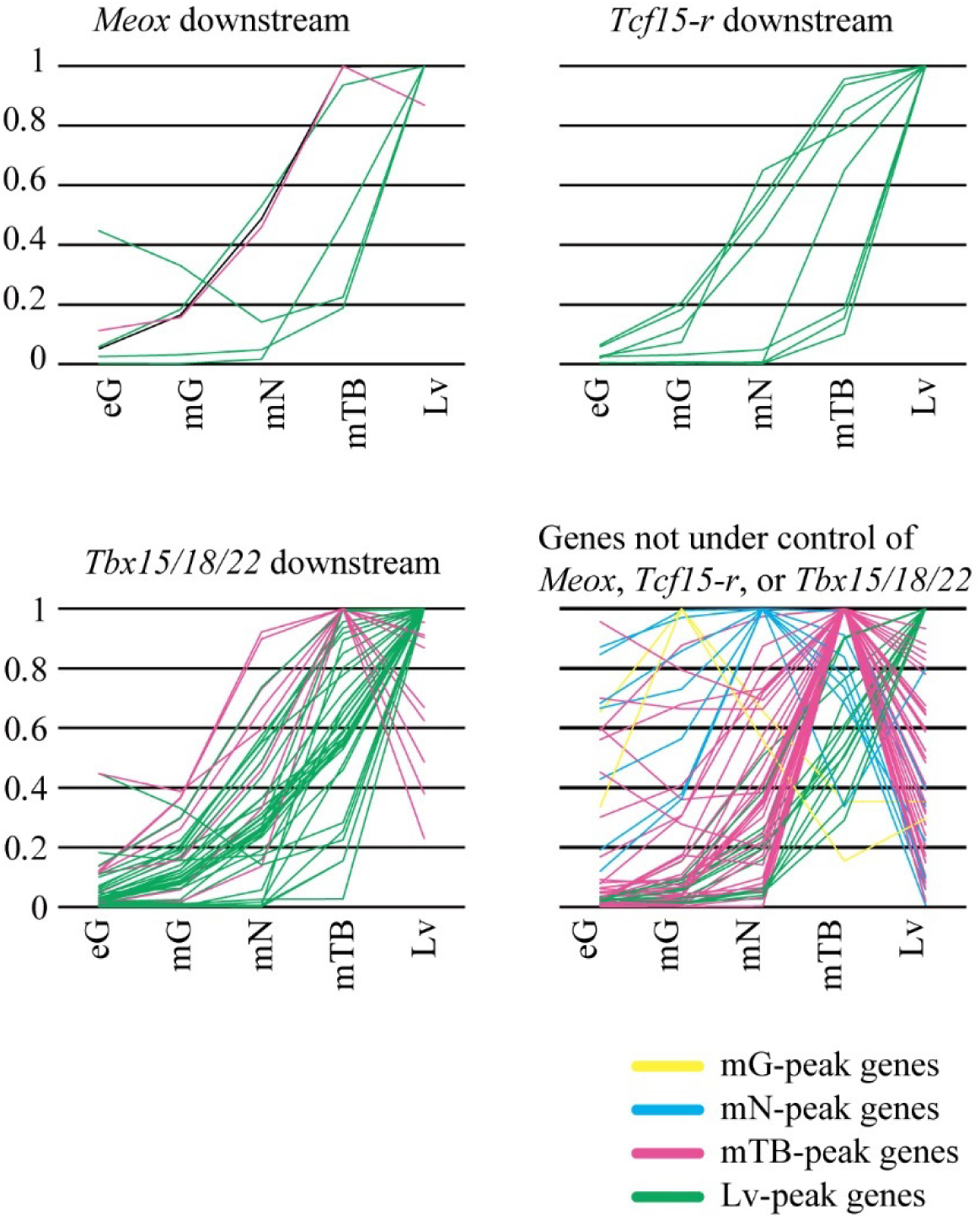
Temporal expression patterns of genes downstream of *Meox*, *Tcf15-r*, and *Tbx15/18/22* and genes that are not under their control. Temporal expression profiles were calculated using publicly available RNA-sequencing data (Brozovic et al., 2018). TPM values were converted to relative values against the highest value for each gene. Genes with peak expression at middle gastrula (mG-peak genes), middle neurula (mN-peak genes), middle tailbud (mTB-peak genes) and larval (Lv-peak genes) stages are shown in yellow, cyan, magenta, and green, respectively. Note that genes that are under control of two or three among *Meox*, *Tcf15-r*, and *Tbx15/18/22* are included in multiple panels.

**Figure S7.**
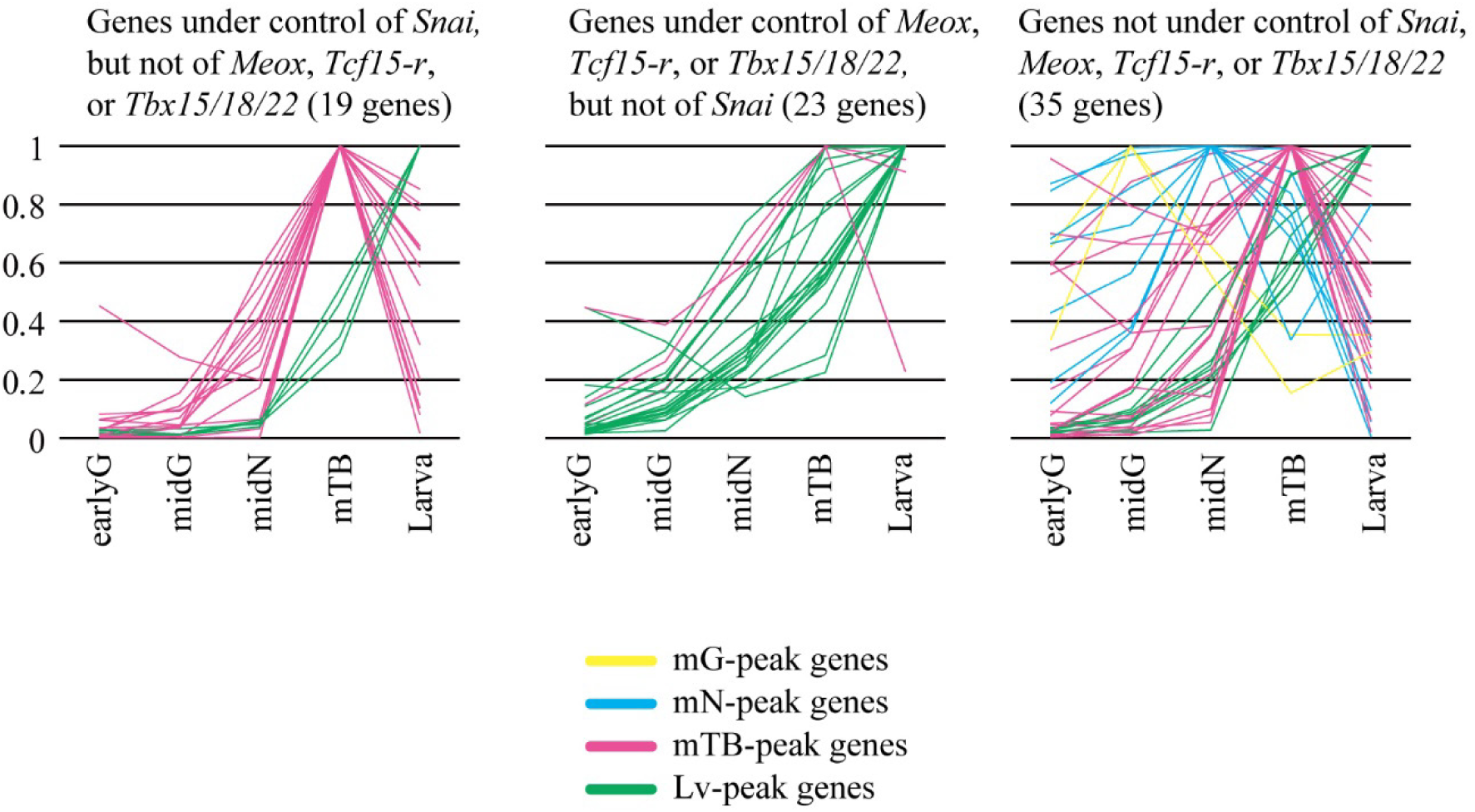
Most genes downstream of *Snai* are highly expressed at the middle tailbud stage. Temporal expression profiles were calculated using publicly available RNA-sequencing data (Brozovic et al., 2018). TPM values were converted to relative values against the highest value for each gene. Genes with peak expression at the middle gastrula (mG-peak genes), middle neurula (mN-peak genes), middle tailbud (mTB-peak genes) and larval (Lv-peak genes) stages are shown in yellow, cyan, magenta, and green, respectively. Three gene populations are shown in different graphs: genes under control of *Snai*, but not of *Meox*, *Tcf15-r*, or *Tbx15/18/22*; genes under control of *Meox, Tcf15-r,* and/or *Tbx15/18/22*; genes not under control of any of them.

**Figure S8.**
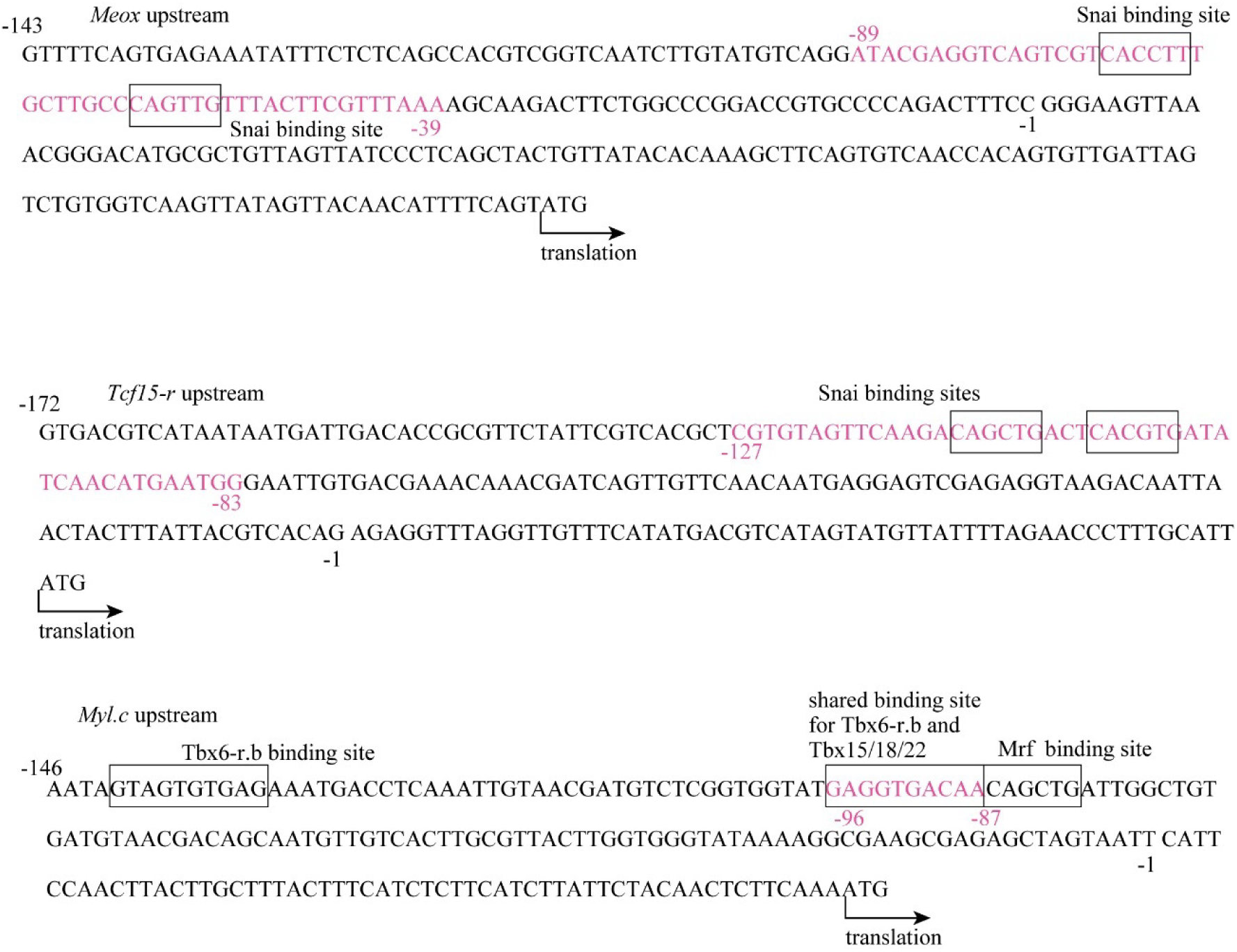
Upstream regulatory regions of *Meox*, *Tcf15-r*, and *Myl.c*. The transcription-factor binding-sites indicated in Figure 6 are enclosed by boxes. Magenta nucleotides in the upstream regions of *Meox* and *Tcf15-r* indicate nucleotide sequences used for the chimera reporter constructs shown in Figure 6FG and gel shift assays shown in Figure 6C. Magenta nucleotide in the upstream region of *Myl.c* indicates the region deleted in Figure 6E.

**Figure S9.**
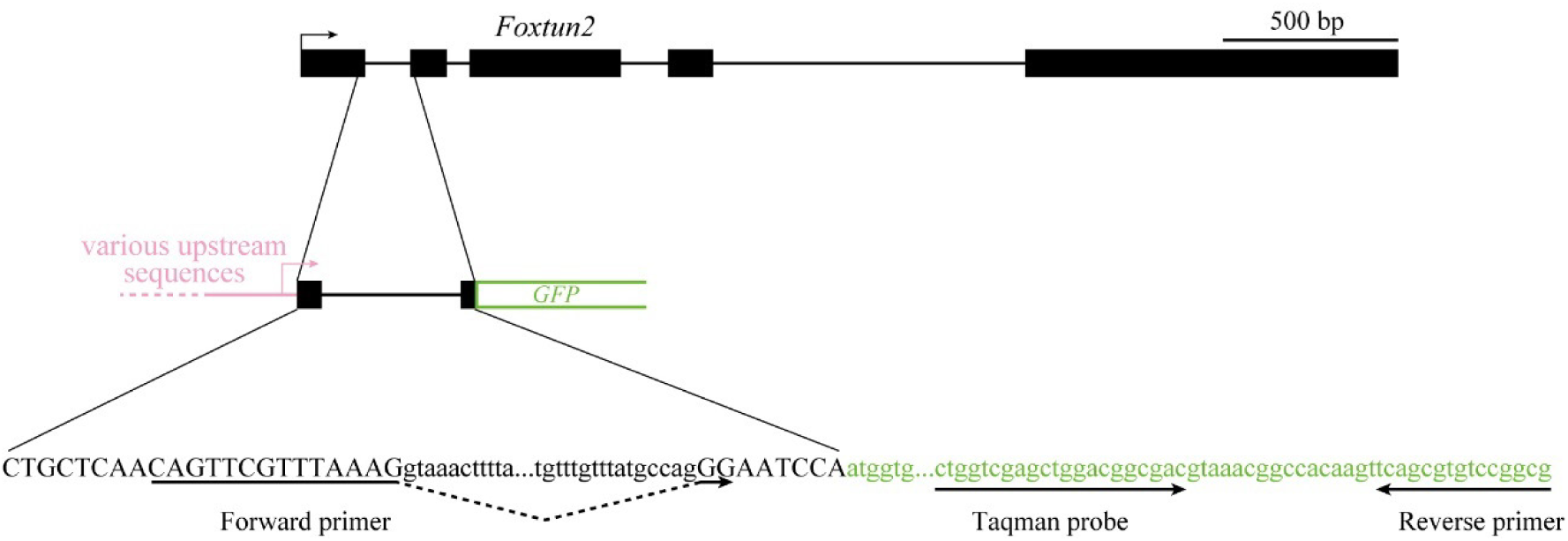
The structure of reporter constructs. Reporter constructs contain part of the first and second exons and the entire first intron of *Foxtun2* (KY21.Chr14.884). The upper panel shows the structure of *Foxtun2* in the genome. Boxes indicate exons, and lines indicate introns. The middle panel shows the structure of reporter constructs. The bottom panel indicates the nucleotide sequence that covers *Foxtun2*-derived region and the initial part of GFP. Irrelevant nucleotides are omitted and shown with dots. Primers and probes used for qPCR are shown with arrows.

## Notes

### Competing Interest Statement

The authors have declared no competing interest.

## References

Altschul, S. F., Gish, W., Miller, W., Myers, E. W. and Lipman, D. J. (1990). Basic local alignment search tool. J Mol Biol 215, 403–410.

Araki, I., Saiga, H., Makabe, K. W. and Satoh, N. (1994). Expression of Amd1, a Gene for a Myod1-Related Factor in the Ascidian Halocynthia-Roretzi. Roux Arch Dev Biol 203, 320–327.

Ashburner, M., Ball, C. A., Blake, J. A., Botstein, D., Butler, H., Cherry, J. M., Davis, A. P., Dolinski, K., Dwight, S. S., Eppig, J. T., et al. (2000). Gene Ontology: tool for the unification of biology. Nature genetics 25, 25–29.

Balagopalan, L., Keller, C. A. and Abmayr, S. M. (2001). Loss-of-function mutations reveal that the Drosophila nautilus gene is not essential for embryonic myogenesis or viability. Dev Biol 231, 374–382.

Brennan, T. J., Edmondson, D. G. and Olson, E. N. (1990). Aberrant Regulation of Myod1 Contributes to the Partially Defective Myogenic Phenotype of Bc3h1 Cells. Journal of Cell Biology 110, 929–937.

Brozovic, M., Dantec, C., Dardaillon, J., Dauga, D., Faure, E., Gineste, M., Louis, A., Naville, M., Nitta, K. R., Piette, J., et al. (2018). ANISEED 2017: extending the integrated ascidian database to the exploration and evolutionary comparison of genome-scale datasets. Nucleic Acids Research 46, D718–D725.

Cao, C., Lemaire, L. A., Wang, W., Yoon, P. H., Choi, Y. A., Parsons, L. R., Matese, J. C., Wang, W., Levine, M. and Chen, K. (2019). Comprehensive single-cell transcriptome lineages of a proto-vertebrate. Nature 571, 349–354.

Chen, L. S., Krause, M., Sepanski, M. and Fire, A. (1994). The Caenorhabditis-Elegans Myod Homolog Hlh-1 Is Essential for Proper Muscle Function and Complete Morphogenesis. Development 120, 1631–1641.

Chopra, V. S. and Levine, M. (2009). Combinatorial patterning mechanisms in the Drosophila embryo. Brief Funct Genomic Proteomic 8, 243–249.

Corbo, J. C., Levine, M. and Zeller, R. W. (1997). Characterization of a notochord-specific enhancer from the Brachyury promoter region of the ascidian, Ciona intestinalis. Development 124, 589–602.

Dehal, P., Satou, Y., Campbell, R. K., Chapman, J., Degnan, B., De Tomaso, A., Davidson, B., Di Gregorio, A., Gelpke, M., Goodstein, D. M., et al. (2002). The draft genome of *Ciona intestinalis*: insights into chordate and vertebrate origins. Science 298, 2157–2167.

Fujiwara, S., Corbo, J. C. and Levine, M. (1998). The snail repressor establishes a muscle/notochord boundary in the Ciona embryo. Development 125, 2511–2520.

Hasty, P., Bradley, A., Morris, J. H., Edmondson, D. G., Venuti, J. M., Olson, E. N. and Klein, W. H. (1993). Muscle Deficiency and Neonatal Death in Mice with a Targeted Mutation in the Myogenin Gene. Nature 364, 501–506.

Hudson, C., Lotito, S. and Yasuo, H. (2007). Sequential and combinatorial inputs from Nodal, Delta2/Notch and FGF/MEK/ERK signalling pathways establish a grid-like organisation of distinct cell identities in the ascidian neural plate. Development 134, 3527-3537.

Ikuta, T. and Saiga, H. (2007). Dynamic change in the expression of developmental genes in the ascidian central nervous system: revisit to the tripartite model and the origin of the midbrain-hindbrain boundary region. Dev Biol 312, 631–643.

Imai, K. S., Hino, K., Yagi, K., Satoh, N. and Satou, Y. (2004). Gene expression profiles of transcription factors and signaling molecules in the ascidian embryo: towards a comprehensive understanding of gene networks. Development 131, 4047–4058.

Imai, K. S., Hudson, C., Oda-Ishii, I., Yasuo, H. and Satou, Y. (2016). Antagonism between β-catenin and Gata.a sequentially segregates the germ layers of ascidian embryos. Development 143, 4167–4172.

Imai, K. S., Levine, M., Satoh, N. and Satou, Y. (2006). Regulatory blueprint for a chordate embryo. Science 312, 1183–1187.

Imai, K. S., Satou, Y. and Satoh, N. (2002). Multiple functions of a Zic-like gene in the differentiation of notochord, central nervous system and muscle in Ciona savignyi embryos. Development 129, 2723–2732.

Imai, K. S., Stolfi, A., Levine, M. and Satou, Y. (2009). Gene regulatory networks underlying the compartmentalization of the Ciona central nervous system. Development 136, 285–293.

Ishida, T. and Satou, Y. (2024). Ascidian embryonic cells with properties of neural-crest cells and neuromesodermal progenitors of vertebrates. Nat Ecol Evol.

Kassar-Duchossoy, L., Gayraud-Morel, B., Gomes, D., Rocancourt, D., Buckingham, M., Shinin, V. and Tajbakhsh, S. (2004). Mrf4 determines skeletal muscle identity in Myf5:Myod double-mutant mice. Nature 431, 466–471.

Katikala, L., Aihara, H., Passamaneck, Y. J., Gazdoiu, S., Jose-Edwards, D. S., Kugler, J. E., Oda-Ishii, I., Imai, J. H., Nibu, Y. and Di Gregorio, A. (2013). Functional Brachyury binding sites establish a temporal read-out of gene expression in the Ciona notochord. PLoS Biol 11, e1001697.

Langmead, B. and Salzberg, S. L. (2012). Fast gapped-read alignment with Bowtie 2. Nature methods 9, 357–U354.

Lassar, A. B., Paterson, B. M. and Weintraub, H. (1986). Transfection of a DNA Locus That Mediates the Conversion of 10t1/2 Fibroblasts to Myoblasts. Cell 47, 649–656.

Long, J. J., Mariossi, A., Cao, C., Mo, Z. Y., Thompson, J. W., Levine, M. S., Satou, Y. and Lemaire, L. A. (2023). Cereblon influences the timing of muscle differentiation in tadpoles. Proceedings of the National Academy of Sciences of the United States of America 120.

Love, M. I., Huber, W. and Anders, S. (2014). Moderated estimation of fold change and dispersion for RNA-seq data with DESeq2. Genome Biology 15.

Meedel, T. H., Chang, P. and Yasuo, H. (2007). Muscle development in Ciona intestinalis requires the b-HLH myogenic regulatory factor gene Ci-MRF. Dev Biol 302, 333–344.

Meedel, T. H., Farmer, S. C. and Lee, J. J. (1997). The single MyoD family gene of Ciona intestinalis encodes two differentially expressed proteins: Implications for the evolution of chordate muscle gene regulation. Development 124, 1711–1721.

Nabeshima, Y., Hanaoka, K., Hayasaka, M., Esumi, E., Li, S. W., Nonaka, I. and Nabeshima, Y. (1993). Myogenin Gene Disruption Results in Perinatal Lethality Because of Severe Muscle Defect. Nature 364, 532–535.

Nishida, H. and Sawada, K. (2001). macho-1 encodes a localized mRNA in ascidian eggs that specifies muscle fate during embryogenesis. Nature 409, 724–729.

Rembold, M., Ciglar, L., Yáñez-Cuna, J. O., Zinzen, R. P., Girardot, C., Jain, A., Welte, M. A., Stark, A., Leptin, M. and Furlong, E. E. M. (2014). A conserved role for Snail as a potentiator of active transcription. Genes & Development 28, 167–181.

Rudnicki, M. A., Schnegelsberg, P. N. J., Stead, R. H., Braun, T., Arnold, H. H. and Jaenisch, R. (1993). Myod or Myf-5 Is Required for the Formation of Skeletal-Muscle. Cell 75, 1351–1359.

Satou, Y., Kawashima, T., Shoguchi, E., Nakayama, A. and Satoh, N. (2005). An integrated database of the ascidian, *Ciona intestinalis*: Towards functional genomics. Zool Sci 22, 837–843.

Satou, Y., Kusakabe, T., Araki, S. and Satoh, N. (1995). Timing of Initiation of Muscle-Specific Gene-Expression in the Ascidian Embryo Precedes That of Developmental Fate Restriction in Lineage Cells. Dev Growth Differ 37, 319–327.

Satou, Y., Nakamura, R., Yu, D., Yoshida, R., Hamada, M., Fujie, M., Hisata, K., Takeda, H. and Satoh, N. (2019). A nearly complete genome of *Ciona intestinalis* type A (*C. robusta*) reveals the contribution of inversion to chromosomal evolution in the genus *Ciona*. Genome biology and evolution 11, 3144–3157.

Satou, Y. and Satoh, N. (1996). Two cis-regulatory elements are essential for the muscle-specific expression of an actin gene in the ascidian embryo. Dev Growth Differ 38, 565–573.

Satou, Y., Takatori, N., Yamada, L., Mochizuki, Y., Hamaguchi, M., Ishikawa, H., Chiba, S., Imai, K., Kano, S., Murakami, S. D., et al. (2001). Gene expression profiles in Ciona intestinalis tailbud embryos. Development 128, 2893–2904.

Satou, Y., Tokuoka, M., Oda-Ishii, I., Tokuhiro, S., Ishida, T., Liu, B. and Iwamura, Y. (2022). A Manually Curated Gene Model Set for an Ascidian, *Ciona robusta* (*Ciona intestinalis* Type A). Zool Sci 39, 253–260.

Satou, Y., Yagi, K., Imai, K. S., Yamada, L., Nishida, H. and Satoh, N. (2002). macho-1-Related genes in Ciona embryos. Dev Genes Evol 212, 87–92.

Simpson, P. (1983). Maternal-Zygotic Gene Interactions during Formation of the Dorsoventral Pattern in Drosophila Embryos. Genetics 105, 615–632.

Takatori, N., Hotta, K., Mochizuki, Y., Satoh, G., Mitani, Y., Satoh, N., Satou, Y. and Takahashi, H. (2004). T-box genes in the ascidian Ciona intestinalis: Characterization of cDNAs and spatial expression. Dev Dyn 230, 743–753.

Tapscott, S. J. (2005). The circuitry of a master switch: Myod and the regulation of skeletal muscle gene transcription. Development 132, 2685–2695.

Thayer, M. J., Tapscott, S. J., Davis, R. L., Wright, W. E., Lassar, A. B. and Weintraub, H. (1989). Positive Auto-Regulation of the Myogenic Determination Gene Myod1. Cell 58, 241–248.

Tokuoka, M., Kobayashi, K., Lemaire, P. and Satou, Y. (2022). Protein kinases and protein phosphatases encoded in the Ciona robusta genome. Genesis 60, e23471.

Tokuoka, M., Kobayashi, K. and Satou, Y. (2018). Distinct regulation of *Snail* in two muscle lineages of the ascidian embryo achieves temporal coordination of muscle development. Development 145, dev163915.

Tokuoka, M., Maeda, K., Kobayashi, K., Mochizuki, A. and Satou, Y. (2021). The gene regulatory system for specifying germ layers in early embryos of the simple chordate. Sci Adv 7, eabf8210.

UniProt-Consortium (2019). UniProt: a worldwide hub of protein knowledge. Nucleic Acids Res 47, D506–D515.

Venuti, J. M., Gan, L., Kozlowski, M. T. and Klein, W. H. (1993). Developmental potential of muscle cell progenitors and the myogenic factor SUM-1 in the sea urchin embryo. Mech Dev 41, 3–14.

Waki, K., Imai, K. S. and Satou, Y. (2015). Genetic pathways for differentiation of the peripheral nervous system in ascidians. Nat Commun 6, 8719.

Yagi, K., Takatori, N., Satou, Y. and Satoh, N. (2005). Ci-Tbx6b and Ci-Tbx6c are key mediators of the maternal effect gene Ci-macho1 in muscle cell differentiation in Ciona intestinalis embryos. Dev Biol 282, 535–549.

Yasuo, H. and Satoh, N. (1993). Function of vertebrate T gene. Nature 364, 582–583.

Yu, D., Iwamura, Y., Satou, Y. and Oda-Ishii, I. (2021). Tbx15/18/22 shares a binding site with Tbx6-r.b to maintain expression of a muscle structural gene in ascidian late embryos. Dev Biol 483, 1–12.

Yu, D., Oda-Ishii, I., Kubo, A. and Satou, Y. (2019). The regulatory pathway from genes directly activated by maternal factors to muscle structural genes in ascidian embryos. Development 146, dev173104.

Yuasa, H. J., Kawamura, K., Yamamoto, H. and Takagi, T. (2002). The structural organization of ascidian Halocynthia roretzi troponin I genes. Journal of biochemistry 132, 135–141.

